# Fractal dimension, occupancy and hotspot analyses of B cell spatial distribution predict clinical outcome in breast cancer

**DOI:** 10.1101/678607

**Authors:** Juliana C. Wortman, Ting-Fang He, Shawn Solomon, Robert Z. Zhang, Anthony Rosario, Roger Wang, Travis Y. Tu, Daniel Schmolze, Yuan Yuan, Susan E. Yost, Xuefei Li, Herbert Levine, Gurinder Atwal, Peter P. Lee, Clare C. Yu

**Affiliations:** Department of Physics and Astronomy, University of California, Irvine, Irvine, CA 92697; Department of Immuno-Oncology, City of Hope Comprehensive Cancer Center and Beckman Research Institute, 1500 East Duarte Road, Duarte, CA 91010; Department of Pathology, City of Hope Comprehensive Cancer Center, 1500 East Duarte Road, Duarte, CA 91010; Department of Medical Oncology and Therapeutics Research, City of Hope Comprehensive Cancer Center, 1500 East Duarte Road, Duarte, CA 91010; Department of Bioengineering and the Center for Theoretical Biological Physics, Rice University, Houston, TX 77030; Department of Bioengineering and Department of Physics, Northeastern University, Boston, MA 02115; Cold Spring Harbor Laboratory, Cold Spring Harbor, NY 11724

**Keywords:** TNBC, TILS, B cells, T cells, tumor microenvironment, spatial distribution, fractal dimensions

## Abstract

While the density of tumor-infiltrating lymphocytes (TILs) is now well known to correlate with clinical outcome, the clinical significance of spatial distribution of TILs is not well characterized. We have developed novel statistical techniques (including fractal dimension differences, a hotspot analysis, a box counting method that we call ‘occupancy’ and a way to normalize cell density that we call ‘thinning’) to analyze the spatial distribution (at different length scales) of various types of TILs in triple negative breast tumors. Consistent with prior reports, the density of CD20^+^ B cells within tumors is not correlated with clinical outcome. However, we found that their spatial distribution differs significantly between good clinical outcome (no recurrence within at least 5 years of diagnosis) and poor clinical outcome (recurrence with 3 years of diagnosis). Furthermore, CD20^+^ B cells are more spatially dispersed in good outcome tumors and are more likely to infiltrate into cancer cell islands. Lastly, we found significant correlation between the spatial distributions of CD20^+^ B cells and CD8^+^ (cytotoxic) T cells (as well as CD3^+^ T cells), regardless of outcome. These results highlight the significance of the spatial distribution of TILs, especially B cells, within tumors.

**Significance Statement:** Immune cells can fight cancer. For example, a patient has a good prognosis when a high density of killer T cells, a type of immune cell that can kill cancer cells, infiltrates into a tumor. However, there is no clear association between prognosis and the density of B cells, another type of immune cell, in a tumor. We developed several statistical techniques to go beyond cell density and look at the spatial distribution, i.e., the pattern or arrangement of immune cells, in tumors that have been removed from patients with triple negative breast cancer. We find that B cells and killer T cells tend to be more spread out in the tumors of patients whose cancer did not recur.

## Introduction

The evolution and progression of cancer is dependent on the tumor microenvironment, which consists of numerous components including cancer cells, collagen fibers, blood vessels, lymph vessels, fibroblasts, and various types of immune cells (Fig. 1A). Recent studies have found that high densities of tumor-infiltrating lymphocytes (TILs) correlate with favorable clinical outcomes in different types of cancer (1-3). For example, higher densities of CD3^+^ and CD8^+^ T cells were associated with a lower rate of recurrence in colorectal carcinoma (4). (CD3^+^ is a marker for all T cells and CD8^+^ is a marker for cytotoxic (killer) T cells.) In the case of patients with triple negative and HER2-positive breast cancers, a higher density of TILs is associated with a better prognosis (5). While the prognostic value of the density of CD3^+^ and CD8^+^ T cells infiltrating tumors is well known, the clinical significance of other types of immune cells, such as B cells, is less clear (6, 7). (B cells can perform a variety of functions including producing antibodies, secreting cytokines (signaling molecules), and presenting antigens to other types of immune cells.)

**Fig. 1:**
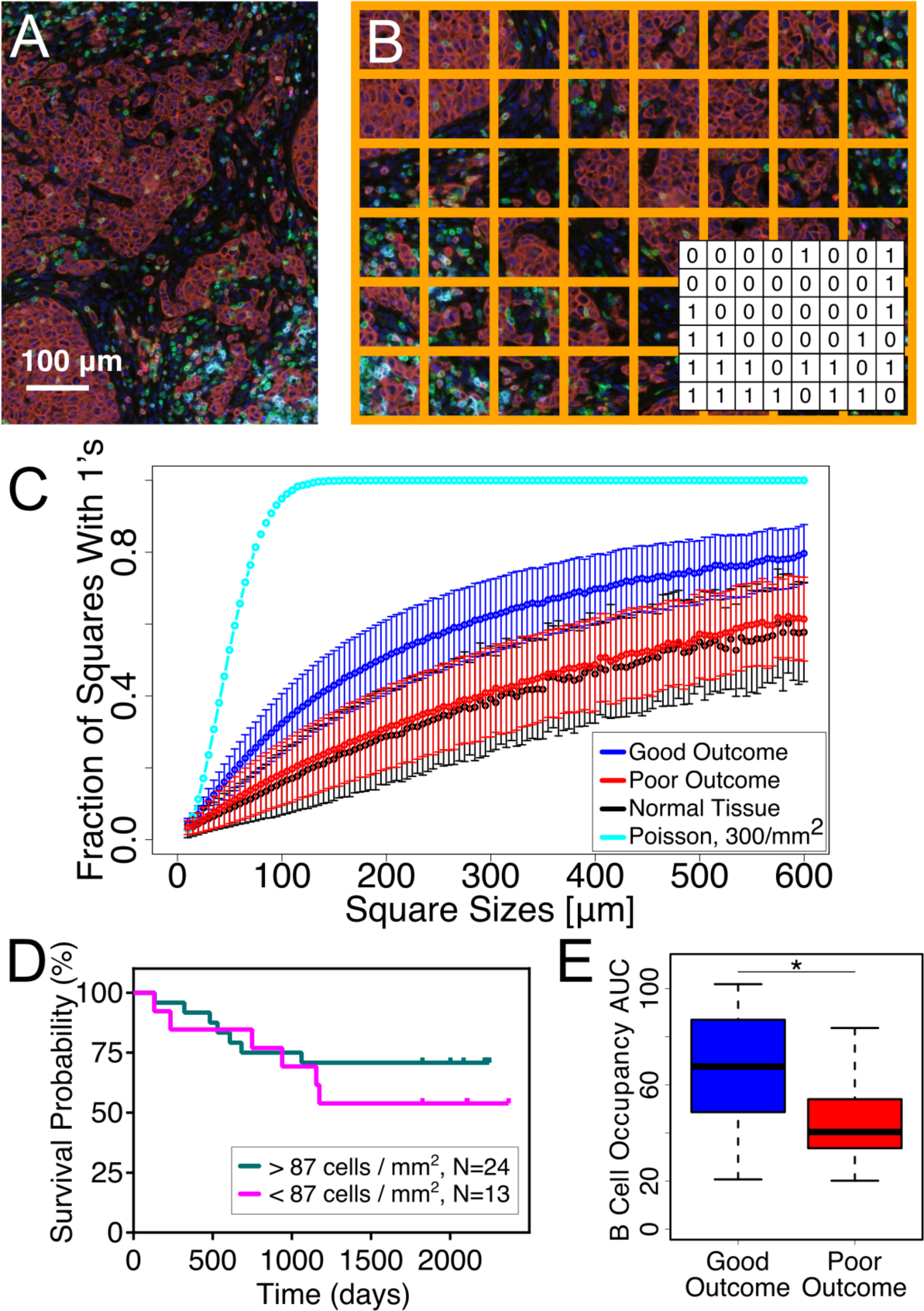
Occupancy. (A) Sample image of TNBC tissue illustrating the heterogeneity of the tumor microenvironment containing various types of immune cells. Color code: T cells (CD3^+^; green), B cells (CD20^+^; cyan), cytotoxic T cells (CD8; bright red), regulatory T cells (FoxP3, magenta), and cancer cells (Pan-cytokeratin cells; dark brownish red). Notice the clusters of cancer cells that we refer to as cancer cell islands (dark brownish red). Scale bar = 100 microns. (B) Grid of squares placed over image. (Inset) 1‘s (0‘s) correspond to yes (no) answers to the question asked of each square, e.g., “Is there at least one B cell in the square?” (C) Plot of CD20^+^ B cell occupancy vs. square size for good clinical outcome (blue), poor clinical outcome (red), normal breast tissue (black) and points randomly distributed according to a uniform Poisson process with a density of 3 × 10^2^ points/mm^2^ (cyan). The error bars indicate 95% confidence intervals. Notice the difference between good and poor clinical outcome. (D) RFS plot showing that CD20^+^ B cell density is not a significant factor in clinical outcome, with a threshold value of 87 cells/mm^2^. (E) Box and whisker plot of CD20^+^ B cell occupancy AUC for entire tumor tissue. The line in the center of each box is the median. Lower and upper part of box are second and third quartiles. Horizontal bars mark maximum and minimum values. This is true of all subsequent box and whisker plots.

If we view the tumor microenvironment in ecological terms, interactions between the different components of this ecosystem depend upon their spatial organization. Regional differences in selective pressures produce microhabitats resulting in phenotypic and genetic heterogeneity (8, 9). Looking at cell densities averaged over the entire tissue overlooks the spatial heterogeneity in the distribution of TILs within the tumor which may be clinically important (10-12). Furthermore, spatial heterogeneity could skew the density measured in tissue microarrays (TMA) which only account for a small region within each tumor - such samples may give a poor estimate of the overall cell density, depending on the location of each TMA sample within the tissue.

This leads to the more general problem of quantifying the spatial distribution (or arrangement) of cells in histology-based images with a single scalar number. Previous efforts to quantify the spatial heterogeneity of the tumor microenvironment include measuring the spatial colocalization of tumor and immune cells (13, 14), locating immune cell clusters or “hotspots” (15, 16), determining the amount of infiltration of lymphocytes into a tumor (17) and using Shannon entropy to quantify the cellular diversity (18). In addition, fractal dimensions (19) have been used to characterize the irregular morphology of tumors (20-22) and tumor-related structures (23-26). (We review these approaches in more detail in the Discussion section.)

It is important to note that these previous approaches produce measures of the spatial distribution at a single length scale. To move beyond this limitation, we have developed unique statistical approaches that use coarse graining to examine the spatial distribution of various types of cells and structures within the tumor microenvironment over a range of length scales. (“Length scale” roughly refers to the magnification factor or zoom level.)

In this paper, we apply these techniques to quantify the spatial distribution of various types of tumor-infiltrating lymphocytes (TILs) in immunohistochemistry-based images of primary tumor tissue from 37 patients with triple negative breast cancer (TNBC) prior to any treatment. According to a recent study of TNBC, 76.2% of recurrences occurred within the first 5 years after surgery, with the median time of recurrence of 2.7 years (27). In this study, we defined patients with no recurrence within 5 years after surgery as good clinical outcome (n=24) and patients who had recurrence within 3 years after surgery as poor clinical outcome (n=13). All the patients were treated with standard chemotherapy; some also had radiotherapy. (See Table S1 for patient clinicopathological characteristics.) As a control, we also analyzed normal breast tissues from 9 patients who underwent breast reduction surgery (age 18-45). As expected, we found that the densities and spatial distributions of CD8^+^ and CD3^+^ T cells differed significantly between patients with good and poor clinical outcome. In contrast, the density of B cells did not correlate with clinical outcome. However, what was surprising was that the spatial distribution of CD20^+^ B cells did differ between good and poor clinical outcome. Furthermore, we found that CD20^+^ B cells were spatially more spread out for good outcome and were more likely to infiltrate into cancer cell islands. Interestingly, we also found that higher CD20+ B cell density specifically within cancer islands was associated with improved prognosis. Lastly, we found significant correlation between the spatial distributions of CD20^+^ B cells and CD8^+^ (as well as CD3^+^) T cells, regardless of clinical outcome.

## Results

### Spatial Distribution of CD20^+^ B Cells in Primary Tumors of TNBC Correlates with clinical response

#### Occupancy

To analyze the spatial distribution of various types of TILs, we first identified the locations of cells in 2D images of archived FFPE tumor tissues from 37 TNBC patients. Each cell type was immunostained and labeled with a different color chromophore. Each image was restricted to tumor-associated regions and excluded necrotic and fibrotic areas determined by a pathologist. (See Methods for more details.) We overlaid each image with a grid of squares as in Fig. 1B. Each square has an area of L^2^. For each square, we then ask a binary (yes-no) question, e.g., “Is there at least one CD20^+^ B cell in the square?” If the answer is yes, we assign a ‘1’ to that square. If the answer is no, we assign a ‘0’ to the square (see inset of Fig. 1B). The *occupancy g* is the fraction of squares with 1‘s, i.e., it is an estimate of the probability that a square will have a 1. To characterize the spatial distribution at different length scales, we varied the size of the squares in the grid and computed the occupancy as a function of L, the length of one side of a square. Note that the occupancy will be affected by the average cell density for questions such as “Is there at least one CD20^+^ B cell in the square?” because the higher the average cell density, the higher the probability that a square is assigned a ‘1’.

To ascertain whether a quantity, such as B cell density, is clinically significant, we calculate the p-value under the null hypothesis and receiver operating characteristic (ROC) area under the curve (AUC), both standard statistical measures of a binary classifier (28). In other words, we assume the clinical outcome is either good or poor. The ROC curve is a parametric curve where the parameter is a cutoff value for the quantity of interest, e.g., B cell density, the x-axis is the false positive rate, and the y-axis is the true positive rate. The ROC AUC varies from 0 to 1. The higher the ROC AUC is, the better the quantity is as a predictor of the outcome. A ROC AUC value of 0.5 means the quantity does no better than random chance.

In Fig. 1C we plot the average occupancy for CD20^+^ B cells, i.e., the fraction of boxes with 1‘s, as a function of the square size L. Notice that the B cells are not randomly distributed. The occupancy for points randomly placed according to a uniform Poisson process is shown as a solid cyan line. The occupancy of CD20^+^ B cells in normal breast tissue (black) is compared to that of CD20^+^ B cells in tumor from good outcome patients (blue) and poor outcome patients (red). Furthermore, notice the difference in occupancy between patients with good and poor clinical outcome. To some extent this reflects the minor difference in B cell density. The average CD20^+^ B cell density in the tumor is slightly higher in patients with good outcome (3.0 × 10^2^/mm^2^) than in those with poor outcome (2.3× 10^2^/mm^2^). However, the CD20^+^ B cell density is not correlated with outcome (p=0.5, ROC AUC=0.6) as is clear from the relapse free survival (RFS) curves in Fig. 1D. On the other hand, the area under the occupancy curve (occupancy AUC) is correlated with outcome (p=0.01, ROC AUC=0.75) as shown in the box and whisker plot in Fig. 1E. (Analogous plots for CD8^+^, CD3^+^, CD4^+^Foxp3^-^, and Foxp3^+^ T cells are shown in the supplement (Figs. S1 and S2). CD4^+^Foxp3^-^ are markers for conventional helper T (Th) cells which can secrete cytokines and activate B cells and cytotoxic T cells. Foxp3^+^ is a marker for regulatory T (Treg) cells which tend to suppress immune response. These plots show that cell density and occupancy AUC is clinically significant for CD8^+^ and CD3^+^ T cells but not for Th and Treg cells.)

#### Thinning

As we mentioned above, occupancy tends to increase with the average cell density. One way to remove the effect of density on the occupancy is to reduce, or thin, the density by randomly eliminating CD20^+^ B cells in the various images until the densities in all the tissue images have the same value as the image with the lowest density. Thinning is explained in greater detail in the supplement where there are plots of thinned occupancy vs. L for CD20^+^ B cells and CD8^+^, CD3^+^, CD4^+^Foxp3^-^ and Foxp3^+^ T cells (Fig. S3). These thinned plots more clearly show that different clinical outcomes have different spatial distributions of these types of cells (Figs. S3B-F). In particular, the (unthinned) occupancy vs. L for CD8^+^ T (and CD3^+^ T) cells differs between good and poor outcome (see Figs. S2A and S2B). In part, this occupancy difference is due to the outcome difference in mean density of CD8^+^ T (and CD3^+^) cells. After thinning, the difference in occupancy between good and poor outcome is more evident for CD20^+^ B cells and the occupancy AUC is much more clinically significant than before thinning (compare Figs. 1C and 1E with Figs. S3B and S3G). This indicates that the spatial distribution, rather than the density, of CD20^+^ B cells differs significantly between good and poor outcomes. In contrast, both the thinned and unthinned occupancy plots for Th and Treg cells do not show a clinical difference (see Figs. S2C-D, S2G-H, S3E-F, S3J-K). Post thinning, the occupancy AUC is barely clinically significant for CD3^+^ T cells (Figs. S3D and S3I) while the occupancy AUC becomes not clinically significant for CD8^+^ T cells (Figs. S3C and S3H).

#### Fractal Dimension (FD)

Another way to characterize the number of boxes with 1‘s and hence, the spatial distribution of cells, is with fractal dimension. Although a number of papers have used fractal dimension to analyze morphologies associated with tumors (20-26), we are using it to quantify the spatial distribution of individual immune cells. While there are a number of different ways to define the fractal dimension, we chose to use a variation of the box counting method (29). As a way of introducing fractal dimension, consider the following simple question: “How many square tiles do you need to tile the floor in a square room?” The answer depends on the size of the tiles, each of which has area L x L. The number of tiles will be proportional to 1/L^2^, where the exponent “2” is the result of the floor being two-dimensional. Now consider a 2D image. As before, we imagine covering the image with a grid of squares as described above, asking a yes-no question of each square, putting a ‘1’ in the square if the answer is yes and a ‘0’ if the answer is no. Let each square have size L x L. The number n(L) of squares with ‘1’ will be proportional to (1/L^δ^) where δ is an exponent less than or equal to the dimension of the image, i.e., 2. In fact, δ is one type of fractal dimension. If the system is self-similar and fractal over a range of length scales L, n(L) should follow a power law: n(L) ∼ (1/L^δ^) and the exponent δ should stay the same as we vary L. So we plot log [n(L)] versus log[A/L] where A is a constant. Then we use linear regression to fit a line through the points. We define the fractal dimension *s* to be the slope of the line: *s* = -d[log n(L)]/d[log L]. (Note that unlike the more common definition of the box-counting fractal dimension, we do not take the limit L**→**0, because we are interested in the distribution of individual cells at different length scales.) If the spatial distribution is truly self-similar, then the plot of log [n(L)] versus log[A/L] will be a straight line and the slope *s* will be independent of L. However, we find that the plots curve so that the slope *s*(L) depends on L. So we use a different variable, *s*(L), rather than δ, since n(L) typically does not follow a simple power law as L is varied. In practice, we find *s*(L) from the slope a line drawn using linear regression of a least squares fit. In applying our technique, the fractal dimension over a range of values of L is determined for each patient and then averaged over all patients with a given clinical outcome.

In Fig. 2A we plot ln(n(L)) versus ln(1/L) for CD20^+^ B cells. At small (10-40 µm) and large (200-600 µm) length scales, we fit the data to straight lines. The slopes of the lines on this log-log plot correspond to the fractal dimension (FD). The closer the fractal dimension is to two, i.e., the larger the fractal dimension is, the more spatially uniform and area-filling the spatial distribution of cells is. Notice that, as expected, randomly placed points (uniform Poisson process) are two-dimensional at length scales that are large compared to the average separation of points.

**Fig. 2:**
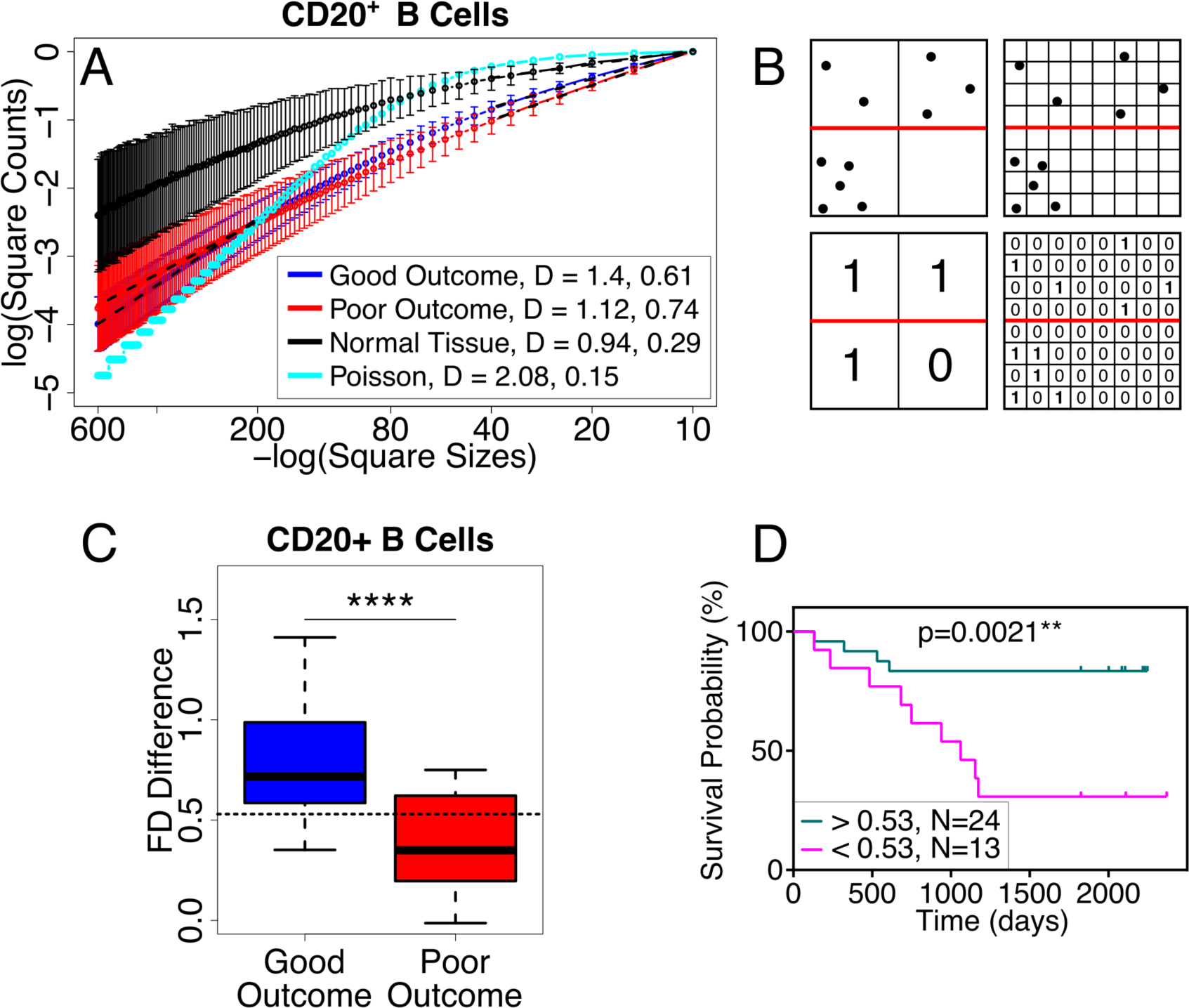
Fractal Dimension and FD difference. (A) Log-log plot of the number of squares with at least one CD20^+^ B cell vs. the inverse box size. (Logarithms are base e.) At long length scales (200-600 microns on the left side of plot), the mean fractal dimension *s* (slope) is 1.4 for good outcome (blue), 1.12 for poor outcome (red), 0.94 for normal tissue (black) and 2.08 for Poisson (cyan). The p value for good vs. poor outcome is 0.02 at long length scales. At short length scales (10-40 microns on the right side of the plot), the mean fractal dimension is 0.61 for good outcome, 0.74 for poor outcome, 0.29 for normal tissue and 0.15 for Poisson. The p value for good vs. poor outcome is 0.1 at short length scales. Black dashed lines show the least squares linear regression fit at long and short length scales. Because different images had different numbers of squares with tissue (cells of any type), we normalized the number n(L) of boxes with 1‘s by the total number N(L) of boxes with cells in computing the fractal dimension. Thus the y-axis values are negative. The error bars correspond to 95% confidence intervals. The slope used to find the fractal dimension is from a least squares fit made using linear regression. (B) Cartoon illustrating how FD difference can tell the difference between spread out cells and clustered cells. The upper halves (above the red lines) of the 2 images show points that are spread out while the lower halves show clustered points. At long length scales (big boxes) the FD is 2 in the upper half but not in the lower half. At small length scales (small boxes), there is the same number of boxes with points and hence the same fractal dimension. (C) Box and whisker plot showing that the FD difference for CD20^+^ B cells is clinically significant (p = 9 × 10^-5^, ROC AUC = 0.86). (D) RFS plot confirms that the FD difference is clinically significant for CD20^+^ B cells (threshold value = 0.53). For all FD difference figures in the main text, the large length scale range is 200-600 microns and the small length scale range is 10-40 microns. Fractal dimension difference and hotspot analysis indicate that CD20^+^ B cells are spatially more spread out in good clinical outcome and more clustered in poor clinical outcome.

#### Difference in fractal dimension

The fractal dimension is larger for good clinical outcome than for poor outcome at large, but this trend is reversed at small length scales. It is useful to look at the difference Δ*s* in fractal dimension between large and small length scales: Δ*s* = *s*_Large_ – *s*_small_, where *s*_Large_ is the fractal dimension at large length scales and *s*_small_ is the fractal dimension at small length scales. The small and large length scales should roughly bracket the typical, or median, nearest neighbor distance between cells of the same type which, in this case, are CD20^+^ B cells. In all the cases we have examined, Δ*s* > 0. If Δ*s* is large, it means that the cells are more dispersed, i.e., more spatially spread out because they appear more two-dimensional at large length scales and more zero-dimensional (point-like) at small length scales (see Fig. 2B). If Δ*s* = 0, the fractal dimension does not change with length scale and the system is self-similar. If Δ*s* is small, then the system is closer to being fractal and the cells are more clustered (see Fig. 2B).

The fractal dimension difference Δ*s* for CD20^+^ B cells is plotted in Fig. 2C where we see that Δ*s* is large (Δ*s* = 0.78) for good outcome, indicating that the B cells are spread out, and small (Δ*s* = 0.38) for poor outcome, indicating that the B cells are somewhat clustered. Furthermore, Fig. 2C shows that the FD difference is a good predictor of outcome (p = 9 × 10^-5^, ROC AUC = 0.86) which is further supported by the RFS plot in Fig. 2D. For CD20^+^ B cells, Δ*s* is not statistically correlated with B cell density (p=0.052, r=0.3), indicating that spatial distribution of B cells is an independent predictor for clinical outcome (Analogous plots of Δ*s* for CD8^+^, CD3^+^, CD4^+^Foxp3^-^, and Foxp3^+^ T cells are shown in the supplement (Figs. S4). Δ*s* is correlated with outcome for CD8^+^ T cells, but not for CD3^+^ T, CD4^+^ Foxp3^-^ T, and Foxp3^+^ T cells. For CD8^+^ T cells, Δ*s* is larger for good outcome, indicating that the CD8^+^ T cells are more spread out in this case. For both CD8^+^ T and CD3^+^ T cells in general, Δ*s* is correlated with their respective cell density. For Th and Treg cells, Δ*s* is not correlated with their respective cell density. Fig. S5 shows the effect of thinning on the FD difference for B cells and various types of T cells. Only thinned CD20^+^ B cells and CD3^+^ T cells have clinically significant FD differences.)

#### Hotspot analysis

Further evidence that the spatial distribution of CD20^+^ B cells differs between good and poor clinical outcome comes from the fraction of the image area where there are ‘hotspots’, i.e., where the density of CD20^+^ B cells is greater than the average value for that image. To do this analysis, each CD20^+^ B cell is represented by a two-dimensional Gaussian distribution that represents the local density due to that cell. The width of the Gaussian is 2σ, i.e., σ^2^ is the dispersion or variance of the Gaussian. We then lay a square lattice of points (with lattice constant *a*) over the image and add the Gaussian weights at each lattice point (see Fig. 3A). The resulting sum is the local CD20^+^ B cell density. We then average the densities over the entire lattice of points of each sample and calculate the fraction of lattice points greater than the average for that image. We refer to this as the fraction of the (image) area with hotspots. Note that this fraction is independent of the value of the average B cell density. In other words, since the hotspots are measured relative to the average density in each particular image, the results are not directly affected by the variation from image to image in the average CD20^+^ B cell density. We then average over images and vary σ.

**Fig. 3:**
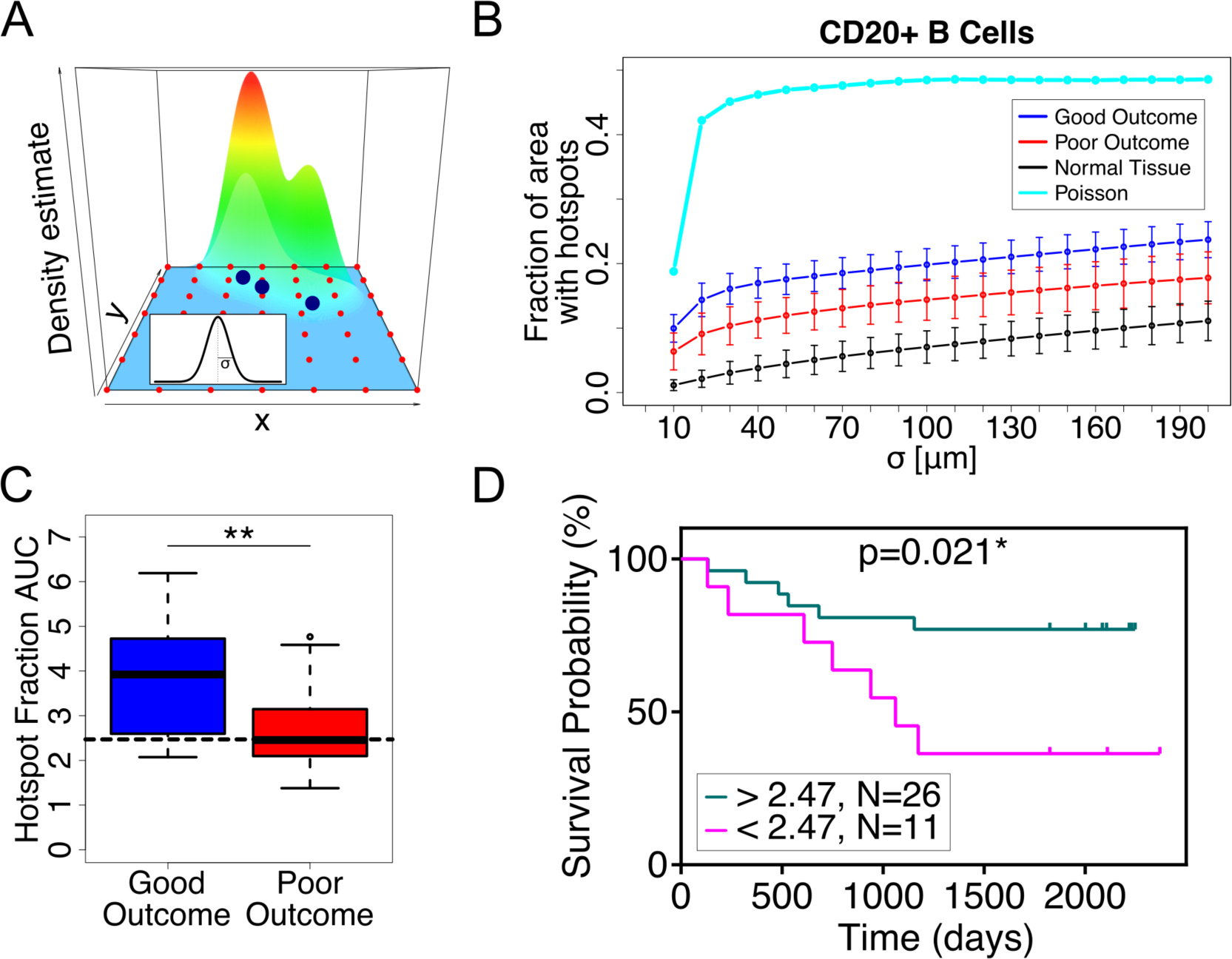
Hotspot analysis. (A) Cartoon illustrating hotspot analysis. Three large blue points indicate the locations of three cells. Each cell‘s contribution to the local density is represented by a Gaussian distribution which is shown in the inset. The mountain over the three cells represents the sum of their Gaussian weights, i.e., the local cell density. The small red points are the grid points where the Gaussian weights are summed. Inset shows a Gaussian curve with width 2σ. (B) Fraction of area with B cell hotspots vs. σ for good outcome (blue), poor outcome (red), normal tissue (black) and random points (cyan). The error bars correspond to 95% confidence intervals. (C) Box and whisker plot of hotspot AUC for CD20^+^ B cells. The larger value of the hotspot AUC for good outcome indicates that the CD20^+^ B cells are more spread out compared to the more clustered CD20^+^ B cells in poor outcome. (D) RFS plot for B cell hotspots shows that the hotspot AUC is mildly significant clinically. The cutoff value is 2.47.

Fig. 3B shows the average fraction of points (area) with hotspots vs. σ. Notice that for points randomly distributed according to a uniform Poisson process, the fraction of area goes to 0.5 for large σ as one would expect. For random points, half the area is above average in density and half is below average. Fig. 3B shows that the average fraction of area occupied by hotspots is greater for good outcome, indicating again that the CD20^+^ B cells are more spread out for good outcome than for poor outcome. The area under the hotspot curve is clinically significant (p=0.008, ROC AUC = 0.77) as is shown in Figs. 3C and 3D. Analogous plots for CD8^+^ T, CD3^+^ T, CD4^+^ Foxp3^-^ T, and Foxp3^+^ T cell hotspots are shown in Fig. S6 which shows that the hotspot AUC is clinically significant for CD8^+^ and CD3^+^ T cells, but not for Th cells. The box and whisker plot for hotspot AUC for Tregs shows no significant difference for good and poor outcome (Fig. S6H), but the RFS plot shows a mildly significant difference. Note that in general, the patient cohorts of the high and low curves of the RFS plots are a mixture of the patients that we have defined as good and poor clinical outcome.

#### Nearest neighbor (NN) distances between cells of a given type

An *a priori* obvious way to quantify how spread out the B cells are in an image would be to measure the mean or median NN distances between B cells (see Fig. S7A). However, most B cells are quite close (5-20 µm) to another B cell, so the NN distance just reflects the (inverse of the) local cell density rather than the spatial dispersion at long length scales. However, as we describe in the supplement, if the B cells (or cells of a given type) are thinned, then the mean or median nearest neighbor distances can be a good measure of how spread out the cells are. Figs. S7B-F show box and whisker plots of the distribution of NN distances after thinning. The difference in the thinned NN distributions between good and poor outcome is most significant for CD8^+^ T, CD3^+^ T and CD20^+^ B cells and less so for Th cells. There is no significant clinically difference of the thinned NN distributions for the Tregs.

### The spatial distribution of CD20^+^ B cells in cancer cell islands is clinically significant

The tumor microenvironment is a heterogeneous mixture of cancer cell islands and stroma (see Fig. 1A). Most immune cells reside in the stroma. However, it is well known that the higher the density of TILs and leukocytes infiltrating into cancer cell islands, the better the prognosis. The “inflamed” tumors were associated with a better response to immunotherapy than the “non-inflamed” and “immune-excluded” tumors (12, 30-33). Similarly, we find that higher densities of B cells as well as CD8^+^ and CD3^+^ T cells (and Th cells and Tregs to a lesser extent) in the cancer cell islands are also correlated with improved prognosis (Fig. S8).

But is the spatial distribution of CD20^+^ B cells in cancer cell islands clinically significant? To answer this, we applied the statistical techniques described above adapted to CD20^+^ B cells in cancer cell islands. To compute the occupancy curve, we impose a grid of squares and ask the yes-no question, “Does the square have at least one CD20 B cell that is in a cancer island?” Thus all the boxes covering stroma will have zeros. The fraction of squares with 1‘s, i.e., the occupancy, is the number of squares with 1‘s divided by the total number of squares covering the image (including the stroma). As Fig. 4A shows, the area under the occupancy curve clearly demonstrates that the spatial distribution of CD20^+^ B cells in cancer cell islands differs significantly between good and poor clinical outcome (p = 9 × 10^-5^, ROC AUC = 0.87). Fig. 4B supports this finding by showing that the fractal dimension (FD) difference for CD20^+^ B cells in cancer cell islands is clinically significant when comparing good and poor outcomes (p=9 × 10^-5^, ROC AUC = 0.85). Furthermore, since the FD difference is higher for good outcome, it indicates that the B cells are more spread out for good outcome.

**Fig. 4:**
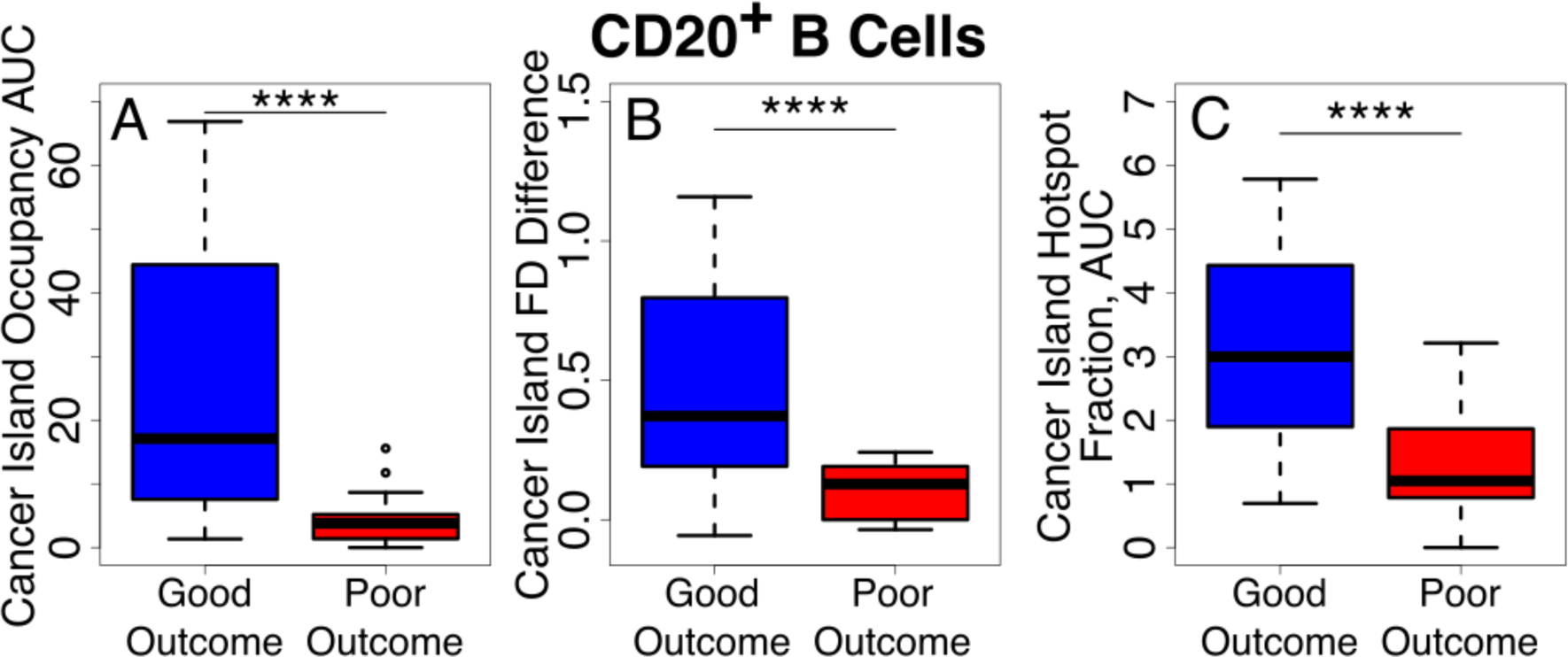
Box and whisker plots of CD20^+^ B cell infiltration into cancer cell islands show strong statistically significant differences between good and poor outcome for: (A) Occupancy AUC (B) FD difference (C) Hotspot AUC.

For the hotspot analysis, only CD20^+^ B cells in cancer cell islands are represented by Gaussians. As the width of the Gaussians increases with σ, the number of grid points outside cancer cell islands with finite CD20^+^ B cell density increases. The average density is the ratio of the sum (over all grid points in the image) of the Gaussian weights divided by the total number of grid points in the entire image instead of only in the cancer cell islands). As before, we plot the fraction of the grid points or area with above average weight (hotspots) vs. σ and compute the area under the curve. As Fig. 4C shows, the area under the hotspot curve (AUC hotspot) is clinically significant (p=2 × 10^-5^, ROC AUC = 0.88). Furthermore, since the AUC hotspot value (3.2) for good outcome is almost three times higher than for poor outcome (1.2), this supports our finding that the B cells are more spread out in cancer cell islands for good outcome. (The supplement shows analogous plots for CD3^+^, CD8^+^, CD4^+^Foxp3^−^, and Foxp3^+^ T cells in Fig. S9 where we see a strong significant difference between good and poor clinical outcome for CD8^+^ and CD3^+^ T cells.)

### The more spatially spread out the CD20^+^ B cells are, the more likely they are to infiltrate the cancer cell islands

We find that compared to poor outcome, the CD20^+^ B cells for good outcome are more spread out as indicated by the higher value of the FD difference (Fig. 2C) and by the higher value of the area under the curve of the fraction of density hotspots vs. σ (Fig. 3C). In addition, good outcome patients have a higher density of B cells that infiltrate cancer cell islands (Fig. 4). This implies that when cells are more spatially dispersed, they are more likely to infiltrate cancer cell islands. We find support for this in Fig. 5A which shows a scatter plot for the correlation between the area under the hotspot curve vs. σ for CD20^+^ B cells in the entire tumor tissue (including stroma and cancer cell islands) and the density of CD20^+^ B cells in cancer cell islands. Each point represents a single patient. The correlation is significant for good outcome (r = 0.54, p=0.007) and for all patients (r = 0.54, p=5 × 10^-4^), but not for poor outcome (r = 0.13, p=0.7). Further support comes from Fig. 5B which shows the correlation between the FD difference for CD20^+^ B cells in the entire tumor tissue and the density of CD20^+^ B cells in cancer cell islands. Again, the correlation is significant for good outcome (r = 0.51, p=0.01) and for all patients (r = 0.56, p=3 × 10^-4^). (Analogous plots for CD8^+^, CD3^+^, CD4^+^Foxp3^−^, and Foxp3^+^ T cells are shown in Fig. S10 where we see significant correlation between the hotspot AUC and the density of cells in cancer islands for CD8^+^, CD3^+^, and CD4^+^Foxp3^−^ T cells.)

**Fig. 5:**
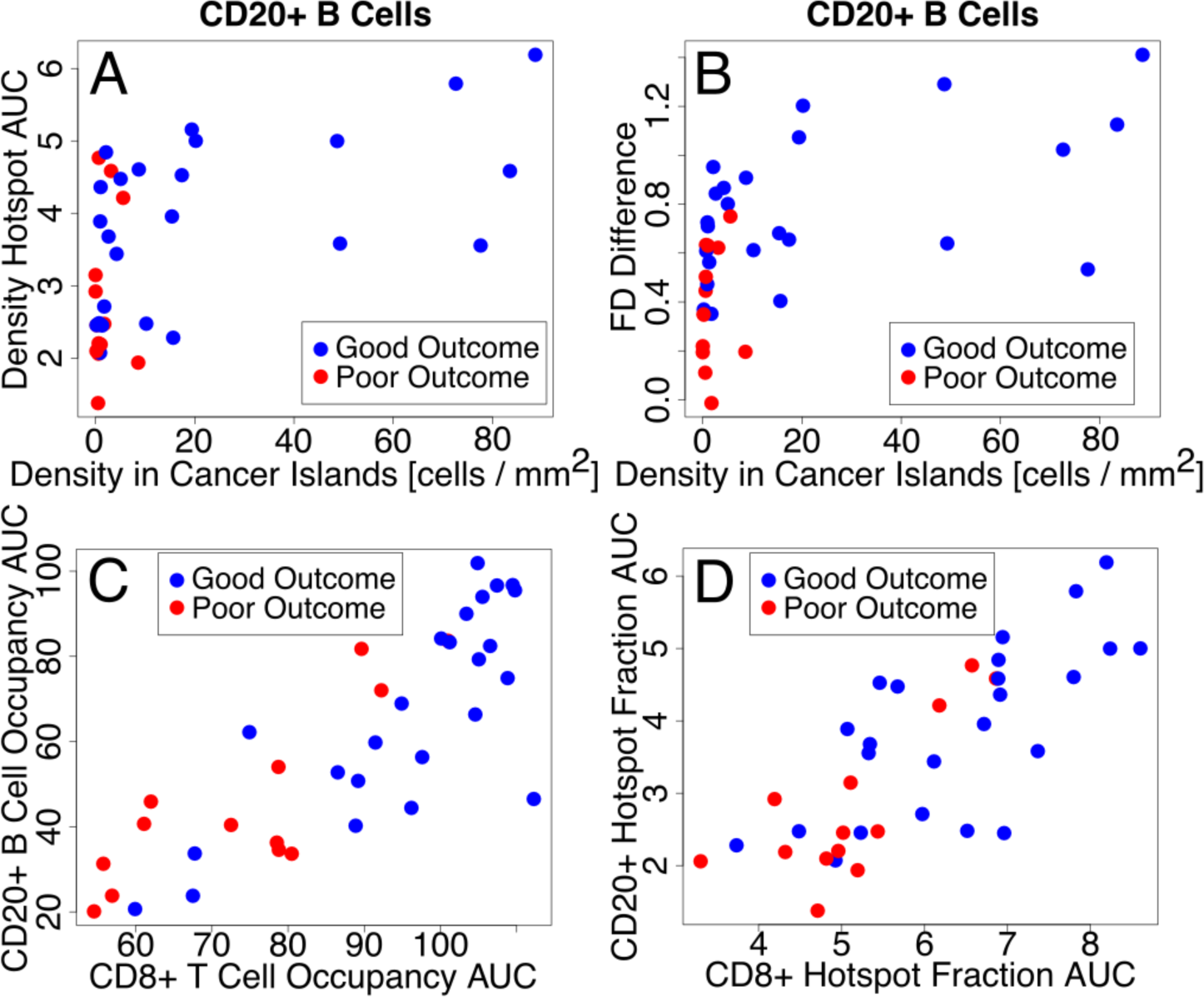
(A-B) Scatter plots showing correlations indicating that the more spread out cells are in the entire tumor tissue, the more likely they are to infiltrate cancer cell islands. (A) B cell hotspot AUC for entire tissue vs. B cell density in cancer cell islands. Pearson r = 0.54, p = 5 × 10^-4^. (B) B cell FD difference ([200-600 µm]-[10-40 µm]) for entire tissue vs. B cell density in cancer cell islands. Pearson r = 0.56, p = 3 × 10^-4^. (C-D) Scatter plots showing the correlation of spatial distributions of B and CD8 T cells in the entire tissue. (C) CD20^+^ B cell occupancy AUC vs. CD8^+^ T cell occupancy AUC in entire tissue. Pearson r = 0.83, p = 3 × 10^-10^. (D) CD20^+^ B cell hotspot AUC vs. CD8^+^ hotspot T cell AUC in entire tissue. Pearson r = 0.78, p = 2 × 10^-8^.

### The spatial distributions of CD20^+^ B cells and CD8^+^ T cells are correlated

The question arises as to whether one could combine the prognostic accuracy of the spatial distribution of CD20^+^ B cells in the entire tumor tissue with the density of CD8^+^ T cells (in the entire tumor tissue) to substantially increase the overall prognostic accuracy. The answer is no because these two quantities are not independent; the spatial distribution of CD20^+^ B cells is correlated with the density as well as the spatial distribution of CD8^+^ T cells. For example, the correlation of the CD20^+^ B cell occupancy AUC with the density of CD8^+^ T cells has a Pearson correlation coefficient r = 0.72 with p = 5 × 10^-7^; and the correlation of the CD20^+^ B cell hotspot AUC with the density of CD8^+^ T cells has r= 0.66 with p = 9 × 10^-6^. The corresponding plots are in Fig. S11 along with plots showing the significant correlation of CD20^+^ B cell occupancy AUC (and CD20^+^ B cell hotspot AUC) with the density of CD3^+^ and CD4^+^Foxp3^-^ T cells. Fig. S11G shows the much weaker correlation of CD20^+^ B cell occupancy AUC with the density of Foxp3^+^ T cells, and Fig. S11H shows the lack of correlation of CD20^+^ B cell hotspot AUC with the density of Foxp3^+^ T cells.

The spatial distribution in the entire tumor tissue of CD20^+^ B cells is also strongly correlated with that of CD8^+^ T cells as shown in Figs. 5C and 5D. Fig. 5C is a scatter plot of the occupancy AUC of CD20^+^ B cells versus that of CD8^+^ T cells (r = 0.83, p = 3 × 10^-10^). Fig. 5D is a scatter plot of the hotspot AUC of CD20^+^ B cells versus that of CD8^+^ T cells (r = 0.78, p = 2 × 10^-8^). (Analogous plots for CD3^+^ T, CD4^+^ Foxp3^-^ T, and Foxp3^+^ T cells are shown in Fig. S12. These plots show that the spatial distribution of CD20^+^ B cells is strongly correlated with that of CD3^+^ T cells and Th cells, but only mildly correlated with that of Tregs.)

## DISCUSSION

Our results show that the spatial distribution of CD20^+^ B cells within the tumor is clinically significant even though their density is not. Furthermore, for good clinical outcome, B cells are more spatially spread throughout the entire tumor and more likely to infiltrate specifically into cancer cell islands compared to poor outcome. We found similar results for CD3^+^ and CD8^+^ T cells (Fig. S10). Finally, the spatial distributions of CD20^+^ B cells and CD8^+^ T cells are correlated, regardless of clinical outcome. These findings raise a number of new questions about the biology of TILs.

One method that we used to characterize the spatial arrangement of cells was fractal dimensions. Self-similarity of the spatial distribution of CD20^+^ B and CD8^+^ T cells is reflected in their fractal dimension at short (10-40 microns) and long (200-600 microns) length scales. But why are the spatial distributions of B and T cells within tumors fractal at all? We speculate that this pattern may arise from the branching trajectories of the B and T cells as they patrol the tissue. Branching structures such as trees and plant roots are self-similar, and hence fractal, because they look the same over a range of length scales, i.e., over a range of magnifications. It may be that the paths that B and T cells travel along have a branching structure because these cells have to go around physical obstacles such as other cells, blood vessels, and collagen fibers. In addition, T cells are known to follow along the outside of blood vessels and collagen fibers (34, 35) which can have a branching architecture.

Second, why do the spatial distributions of B (and T) cells differ between good and poor clinical outcome? Some of the difference in occupancy AUC reflects the difference in B (or T) cell density, but other quantities, such as the FD difference of B cells and the hotspot AUC, that do not depend on density, differ significantly depending on clinical outcome. One could also ask why B (and T) cells are more spread out for good clinical outcome. One possibility is that the difference in spatial distribution reflects differences in the spatial topography (obstacles) of the tumor microenvironment. In adjacent FFPE sections, we performed immunohistochemical staining of collagen I which did not yield differences in the spatial distribution of collagen I between good and poor outcome (data not shown). This indicates that differences in physical topography may not be the explanation. It may be that CD20^+^ B cells, CD8^+^ T cells and CD3^+^ T cells in good outcome patients are more responsive to chemokines secreted by tumor cells and are more successful in finding their cognate antigens within the tumor microenvironment as in a recently published study (36).

Third, why are the spatial distributions of CD20^+^ B cells and CD8^+^ T cells spatially correlated, regardless of clinical outcome? This likely indicates that the motility of both cell types is affected by the same set of factors. We could not determine whether this leads to co-localization of CD20^+^ B cells and CD8^+^ T cells because of the confounding fact that both cell types tend to be sequestered in the stroma, especially for tumors from poor outcome patients.

There have been a number of efforts to quantify spatial heterogeneity of the tumor microenvironment based on comparing populations of cells (13). For example, the Morisita-Horn index was used to quantify the spatial colocalization of tumor and immune cells, and it was found that significant colocalization was associated with a higher disease-specific survival in Her2-positive breast cancers (13, 14). The Getis-Ord analysis (15) was used to locate immune hotspots where the clustering of immune cells was significantly above background. A combined immune-cancer hotspot score was found to be associated with good prognosis in ER-negative breast cancer (16). A quantitative measure of the infiltration of immune cells into a tumor is the intratumor lymphocyte ratio (ITLR) which is defined as the ratio of the number of intratumor lymphocytes to the total number of cancer cells in a histological sample (17). A high ITLR was found to be associated with good disease specific survival in ER-negative/Her2-negative breast cancer (17).

Natrajan *et al.* (18) quantified the spatial heterogeneity in breast tumors with regard to cancer cells, lymphocytes and stromal cells by calculating the Shannon entropy *d*_*i*_ in different regions of the tumor and using Gaussian mixture models to fit the distribution of Shannon entropies. Their ecosystem diversity index (EDI) was the number of Gaussians needed to fit the distribution. They found that high EDI values were associated with high micro-environmental diversity and poor prognosis. Somewhat ironically, with this measure, if most of the regions have high Shannon entropies such that a single Gaussian can be used to fit the distribution, then the EDI is low.

Fractal dimensions (19) have been used to characterize the irregular morphology of tumors (20-22) and vasculature (23, 24) as well as subcellular structures such as mitochondria (25) and nuclei (26). There are numerous ways to calculate fractal dimensions. In the box counting method, the number *N* of squares (each with area L^2^) needed to cover the 2D image of a tumor, say, is proportional to L^-d^, where d is the fractal dimension in the limit that L goes to zero (or a very small value). The more irregular the shape, the higher the fractal dimension is and the poorer the prognosis (20, 21).

Assuming that the structure of tumor tissue is reflected in the arrangement of cancer cell nuclei, Waliszewski *et al*. calculated several different fractal dimensions as well as the Shannon entropy and lacunarity to characterize the spatial distribution of cancer cell nuclei in prostate tumor tissue and compared the results to the corresponding Gleason scores in an attempt to find a more objective way to classify prostate tumor tissue (37, 38).

The statistical techniques that we have developed and presented here to quantify spatial distributions are novel: (a) Unlike previous efforts to analyze spatial distributions, we have examined how the spatial distribution varies with length scale and how that can shed light on whether the cells are clustered or spread out spatially. In analyzing the fractal dimension, we did not take the limit of L going to zero (or a very small value) as is usually done, but found it useful to look at the FD difference between large and small length scales. (b) Our hotspot analysis differs from the Getis-Ord hotspot analysis (13, 15) which requires dividing the image up into regions and depends on the cell counts in neighboring regions. Our hotspot method is completely local in the sense that the cell density at one grid point does not depend on the density at neighboring points. (c) Both occupancy and fractal dimension depend on the number of squares with 1‘s, so there must be a simple relation between the dependence of occupancy on L and fractal dimension d. To find the relation, consider the following. Suppose the total number N(L) of squares covering the image of the tissue goes as (1/L^D^). Then if n(L) ∼ (1/L^δ^), the occupancy g = n(L)/N(L) ∼ L^D-δ^. Note that D need not be equal to 2 since the image of the tissue may be irregular or there may be regions that were not imaged.

On a more general level, these approaches are flexible and can be applied to a broad spectrum of problems. There are straightforward applications of these approaches, e.g., to quantify the spatial distributions of various kinds of cells in metastatic tumors or to ascertain which patients are good candidates for various therapeutic treatments. However, one can go beyond simply the spatial distribution of cells or discrete entities. In determining the occupancy, fractal dimension and the difference in fractal dimensions, we laid down a grid of squares and asked a binary yes-no question of each square. In this paper, we asked questions like “Is there at least one CD20^+^ B cell in this square?” However, in other contexts, one may be interested in other questions such as that of co-localization. For example, one could ask “Does the square have at least one T cell that is within 50 microns of a dendritic cell?” or “Does the square have at least one B cell and one T cell within 25 microns of a blood vessel?” Furthermore, these approaches can have applications to image analysis problems outside of biology, e.g., the distribution of galaxies in astrophysics.

Our results highlight the importance of spatial distribution of TILs within tumors with regard to clinical outcome. These findings raise new questions about the biology and clinical impact of TILs. Of particular interest is the discordant significance of B cell density vs. spatial distribution on outcome of TNBC. The role of B cells within tumors is understudied and unclear, partially due to prior reports that their density within tumors is of unclear significance. Our finding of strong correlation of B cell spatial distribution with clinical outcome in TNBC revives their significance in cancer, and opens up new questions on their functional role in preventing recurrence.

------

## METHODS

### Breast Cancer Tissue Sample Preparation and Analysis

#### Tissue preparation, multi-color antibody staining, and image analysis

Specimens were identified through an IRB-approved protocol via the City of Hope (COH) Biospecimen Repository which is funded in part by the National Cancer Institute. Samples from patients diagnosed with triple negative breast cancer and treated at COH from January 1, 1994 to March 4, 2015 were retrieved. Eligible patients had the following features: stage I-III breast cancer; at least one tumor biospecimen was available from the initial surgical resection or biopsy; clinical outcome data was available for identification of relapse free survival; no prior treatment at the time of surgical biopsy. Archived formalin-fixed paraffin-embedded (FFPE) tumor tissues were sectioned (i.e., 3-5 microns per slide) and on the same slide, multi-color immunostaining was performed including anti-pan cytokeratin (AE1/AE3, Dako), anti-CD8 (SP16, Biocare), anti-CD3 (Polyclonal, Dako), anti-FOXP3 (236A/E7, Biocare) and anti-CD20 (L26, Dako). Samples were further counterstained with DAPI to visualize the nuclei of all cells. Prior to imaging, the tissue sections were coverslipped with ProLong^®^ Gold Antifade mounting media (Cat. # P36930, Life Technologies). All the images were acquired using the Vectra 3.0 Automated Quantitative Pathology Imaging System (Akoya Biosciences) and commercially available software packages (inForm, Akoya Biosciences) were used to identify each cell, define its type (cancer or specific immune), and assign it Cartesian coordinates. Using an automated tissue segmenter algorithm built in inForm®, we further divided the images into areas of cancer islands and stroma based on anti-pan cytokeratin antibody staining.

#### Regions of Interest

A pathologist delineated tumor regions that were deemed representative of the entire tumor in terms of cellularity and the tumor-infiltrating lymphocyte (TIL) distribution, and that were free of artifacts such as tissue folding. Obvious large swaths of necrotic tumor were avoided, as were peri-tumoral lymphoid aggregates not in close proximity to the tumor cells. Given these constraints, regions were chosen to maximize tumor area.

#### Spatial Distribution Analysis of TILs

Using the data spreadsheets with every cell phenotyped and given a set of Cartesian coordinates, we tiled each image with a grid of identical squares. We omitted squares that had no cells, since they lay outside the tissue. However, we included squares that were within the tissue but contained no cells of interest. The length L of one side of a square varies from 10 µm to 600 µm. To determine if a square receives a ‘1’ or a ‘0’, we asked a yes-no question, e.g., “Is there at least one cell of a given phenotype in this square?” The phenotypes of interest for this paper were primarily CD20^+^ B cells, CD3^+^ T cells and CD8^+^ T cells. For comparison we calculated the occupancy, fractal dimension and hotspot density of a uniform random (Poisson) distribution of points. We processed the data using in-house custom developed software (RStudio) and the R packages “spatstat” for point patterns and “EBImage” for raster images (39, 40).

## ACKNOWLEDGEMENTS

We thank Dr. Ching Ouyang for helpful discussions. CCY thanks Dr. Larry Norton for a helpful discussion. This work was supported by Stand Up To Cancer, The V Foundation, and the Breast Cancer Research Foundation. The work of CCY and JCW was also supported in part by the Cure Breast Cancer Foundation. The work of CCY was performed in part at the Aspen Center for Physics, which is supported by National Science Foundation grant PHY-1607611.

