## Supplementary Material for "Fractal dimension, occupancy and hotspot analyses of B cell spatial distribution predict clinical outcome in breast cancer"

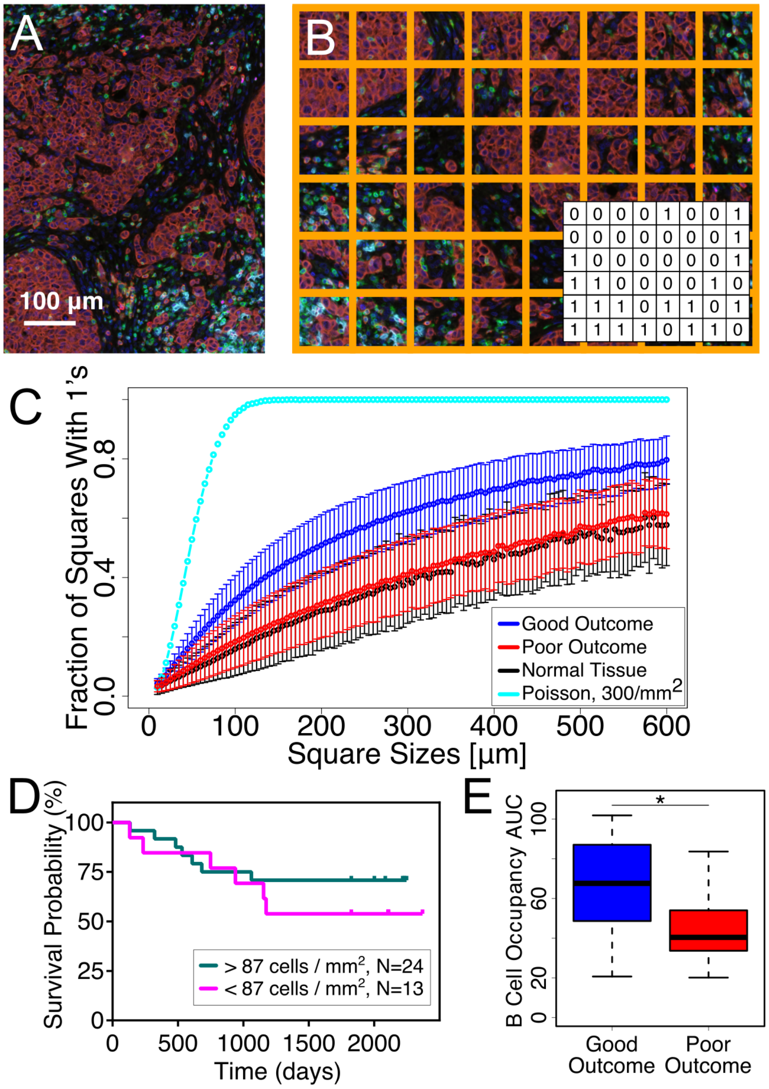

**Fig. 1: Occupancy. (A) Sample image of TNBC tissue illustrating the heterogeneity of the tumor microenvironment containing various types of immune cells. Color code: T cells (CD3$^+$; green), B cells (CD20$^+$; cyan), cytotoxic T cells (CD8; bright red), regulatory T cells (FoxP3, magenta), and cancer cells (Pan-cytokeratin cells; dark brownish red). Notice the clusters of cancer cells that we refer to as cancer cell islands (dark brownish red). Scale bar = 100 microns. (B) Grid of squares placed over image. (Inset) 1's (0's) correspond to yes (no) answers to the question asked of each square, e.g., "Is there at least one B cell in the square?" (C) Plot of CD20$^+$ B cell occupancy vs. square size for good clinical outcome (blue), poor clinical outcome (red), normal breast tissue (black) and points randomly distributed according to a uniform Poisson process with a density of 3 x 10$^2$ points/mm$^2$ (cyan). The error bars indicate 95% confidence intervals. Notice the difference between good and poor clinical outcome. (D) RFS plot showing that CD20$^+$ B cell density is not a significant factor in clinical**

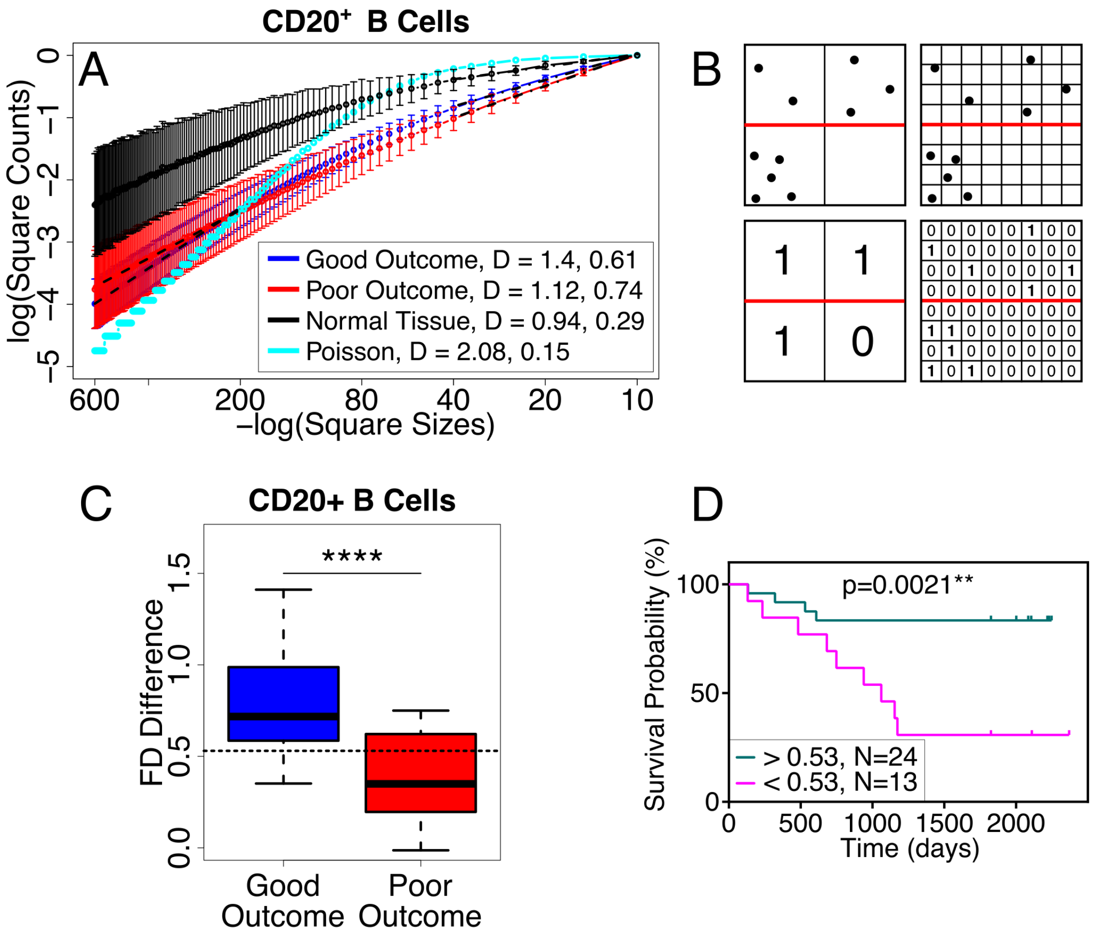

**Fig. 2: Fractal Dimension and FD difference. (A) Log-log plot of the number of squares with at least one CD20+ B cell vs. the inverse box size. (Logarithms are base e.) At long length scales (200-600 microns on the left side of plot), the mean fractal dimension *s* (slope) is 1.4 for good outcome (blue), 1.12 for poor outcome (red), 0.94 for normal tissue (black) and 2.08 for Poisson (cyan). The p value for good vs. poor outcome is 0.02 at long length scales. At short length scales (10-40 microns on the right side of the plot), the mean fractal dimension is 0.61 for good outcome, 0.74 for poor outcome, 0.29 for normal tissue and 0.15 for Poisson. The p value for good vs. poor outcome is 0.1 at short length scales. Black dashed lines show the least squares linear regression fit at long and short length scales. Because different images had different numbers of squares with tissue (cells of any type), we normalized the number n(L) of boxes with 1's by the total number N(L) of boxes with cells in computing the fractal dimension. Thus the y-axis values are negative. The error bars correspond to 95% confidence intervals. The slope used to find the fractal dimension is from a least squares fit made using linear regression. (B) Cartoon illustrating how FD difference can tell the difference between spread out cells and clustered cells. The upper halves (above the red lines) of the 2 images show points that are spread out while the lower halves show clustered points. At long length scales (big boxes) the FD is 2 in the upper half but not in the lower**

**half. At small length scales (small boxes), there is the same number of boxes with points and hence the same fractal dimension. (C) Box and whisker plot showing that the FD difference for $CD20^+$ B cells is clinically significant ($p = 9 \times 10^{-5}$, ROC AUC = 0.86). (D) RFS plot confirms that the FD difference is clinically significant for $CD20^+$ B cells (threshold value = 0.53). For all FD difference figures in the main text, the large length scale range is 200-600 microns and the small length scale range is 10-40 microns.**

**Fractal dimension difference and hotspot analysis indicate that $CD20^+$ B cells are spatially more spread out in good clinical outcome and more clustered in poor clinical outcome.**

*Difference in fractal dimension:* The fractal dimension is larger for good clinical outcome than for poor outcome at large, but this trend is reversed at small length scales. It is useful to look at the difference $\Delta s$ in fractal dimension between large and small length scales: $\Delta s = s_{\text{Large}} - s_{\text{small}}$, where $s_{\text{Large}}$ is the fractal dimension at large length scales and $s_{\text{small}}$ is the fractal dimension at small length scales. The small and large length scales should roughly bracket the typical, or median, nearest neighbor distance between cells of the same type which, in this case, are $CD20^+$ B cells. In all the cases we have examined, $\Delta s > 0$. If $\Delta s$ is large, it means that the cells are more dispersed, i.e., more spatially spread out because they appear more two-dimensional at large length scales and more zero-dimensional (point-like) at small length scales (see Fig. 2B). If $\Delta s = 0$, the fractal dimension does not change with length scale and the system is self-similar. If $\Delta s$ is small, then the system is closer to being fractal and the cells are more clustered (see Fig. 2B).

The fractal dimension difference $\Delta s$ for $CD20^+$ B cells is plotted in Fig. 2C where we see that $\Delta s$ is large ($\Delta s = 0.78$) for good outcome, indicating that the B cells are spread out, and small ($\Delta s = 0.38$) for poor outcome, indicating that the B cells are somewhat clustered. Furthermore, Fig. 2C shows that the FD difference is a good predictor of outcome ($p = 9 \times 10^{-5}$, ROC AUC = 0.86) which is further supported by the RFS plot in Fig. 2D. For $CD20^+$ B cells, $\Delta s$ is not statistically correlated with B cell density (p=0.052, r=0.3), indicating that spatial distribution of B cells is an independent predictor for clinical outcome (Analogous plots of $\Delta s$ for $CD8^+$, $CD3^+$, $CD4^+Foxp3^-$, and $Foxp3^+$ T cells are shown in the supplement (Figs. S4). $\Delta s$ is correlated with outcome for $CD8^+$ T cells, but not for $CD3^+$ T, $CD4^+$ $Foxp3^-$ T, and $Foxp3^+$ T cells. For $CD8^+$ T cells, $\Delta s$ is larger for good outcome, indicating that the $CD8^+$ T cells are more spread out in this case. For both $CD8^+$ T and $CD3^+$ T cells in general, $\Delta s$ is correlated with their respective cell density. For Th and Treg cells, $\Delta s$ is not correlated with their respective cell density. Fig. S5 shows the effect of thinning on the FD difference for B cells and various types of T cells. Only thinned $CD20^+$ B cells and $CD3^+$ T cells have clinically significant FD differences.)

*Hotspot analysis*: Further evidence that the spatial distribution of $CD20^+$ B cells differs between good and poor clinical outcome comes from the fraction of the image area where there are 'hotspots', i.e., where the density of $CD20^+$ B cells is greater than the average value for that image. To do this analysis, each $CD20^+$ B cell is represented by a two-dimensional Gaussian distribution that represents the local density due to that cell. The width of the Gaussian is $2\sigma$, i.e., $\sigma^2$ is the dispersion or variance of the Gaussian. We then lay a square lattice of points (with lattice constant $a$) over the image and add the Gaussian weights at each lattice point (see Fig. 3A). The resulting sum is the local $CD20^+$ B cell density. We then average the densities over the entire lattice of points of each sample and calculate the fraction of lattice points greater than the average for that image. We refer to this as the fraction of the (image) area with hotspots. Note that this fraction is independent of the value of the average B cell density. In other words, since the hotspots are measured relative to the average density in each particular image, the results are not directly affected by the variation from image to image in the average $CD20^+$ B cell density. We then average over images and vary $\sigma$.

Fig. 3B shows the average fraction of points (area) with hotspots vs. $\sigma$. Notice that for points randomly distributed according to a uniform Poisson process, the fraction of area goes to 0.5 for large $\sigma$ as one would expect. For random points, half the area is above average in density and half is below average. Fig. 3B shows that the average fraction of area occupied by hotspots is greater for good outcome, indicating again that the $CD20^+$ B cells are more spread out for good outcome than for poor outcome. The area under the hotspot curve is clinically significant ($p=0.008$, ROC AUC = 0.77) as is shown in Figs. 3C and 3D. Analogous plots for $CD8^+$ T, $CD3^+$ T, $CD4^+$ $Foxp3^-$ T, and $Foxp3^+$ T cell hotspots are shown in Fig. S6 which shows that the hotspot AUC is clinically significant for $CD8^+$ and $CD3^+$ T cells, but not for Th cells. The box and whisker plot for hotspot AUC for Tregs shows no significant difference for good and poor outcome (Fig. S6H), but the RFS plot shows a mildly significant difference. Note that in general, the patient cohorts of the high and low curves of the RFS plots are a mixture of the patients that we have defined as good and poor clinical outcome.

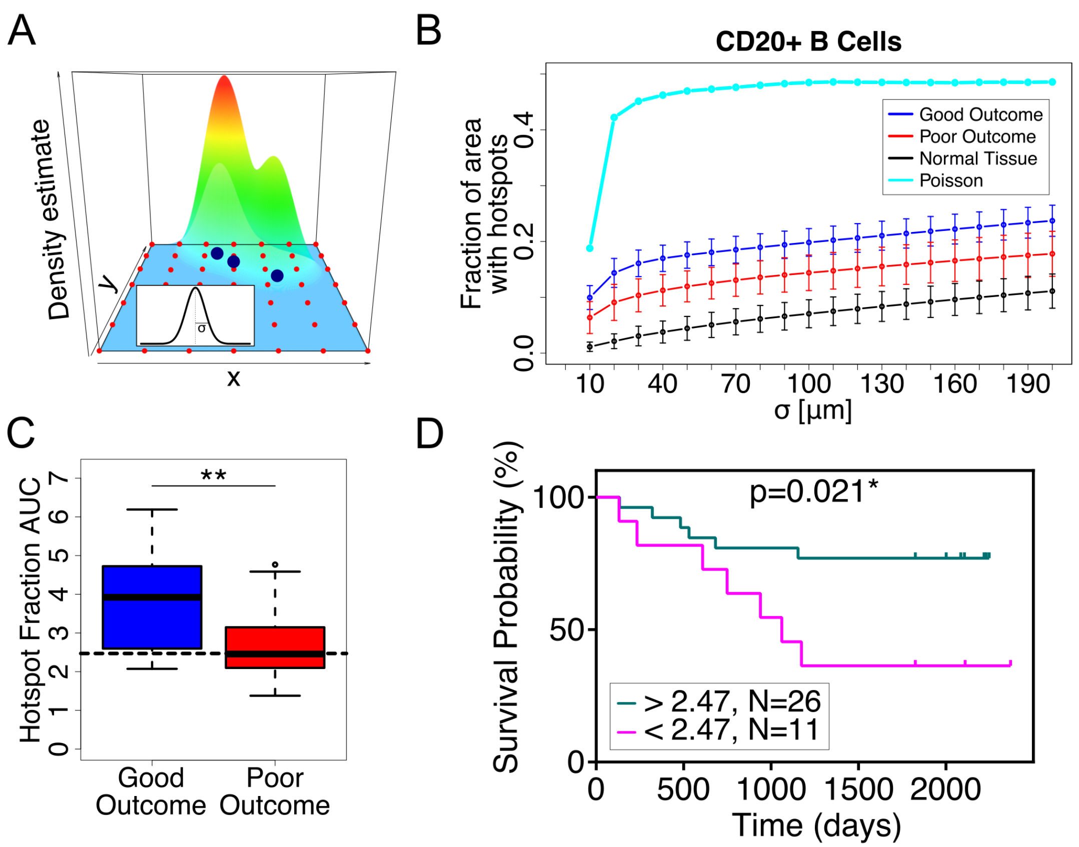

**Fig. 3: Hotspot analysis. (A) Cartoon illustrating hotspot analysis. Three large blue points indicate the locations of three cells. Each cell's contribution to the local density is represented by a Gaussian distribution which is shown in the inset. The mountain over the three cells represents the sum of their Gaussian weights, i.e., the local cell density. The small red points are the grid points where the Gaussian weights are summed. Inset shows a Gaussian curve with width 2σ. (B) Fraction of area with B cell hotspots vs. σ for good outcome (blue), poor outcome (red), normal tissue (black) and random points (cyan). The error bars correspond to 95% confidence intervals. (C) Box and whisker plot of hotspot AUC for CD20$^+$ B cells. The larger value of the hotspot AUC for good outcome indicates that the CD20$^+$ B cells are more spread out compared to the more clustered CD20$^+$ B cells in poor outcome. (D) RFS plot for B cell hotspots shows that the hotspot AUC is mildly significant clinically. The cutoff value is 2.47.**

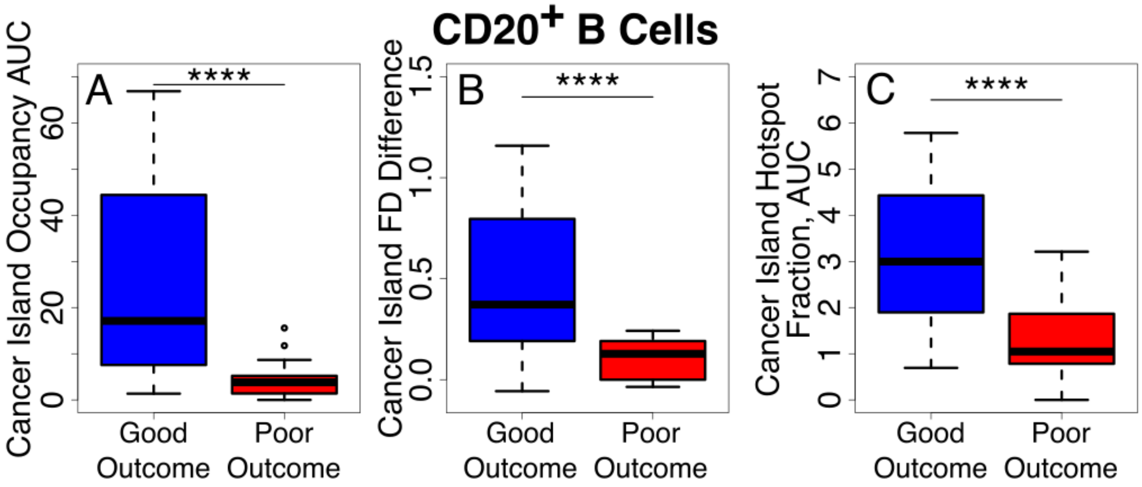

**Fig. 4: Box and whisker plots of CD20$^+$ B cell infiltration into cancer cell islands show strong statistically significant differences between good and poor outcome for: (A) Occupancy AUC (B) FD difference (C) Hotspot AUC.**

**The more spatially spread out the CD20$^+$ B cells are, the more likely they are to infiltrate the cancer cell islands.** We find that compared to poor outcome, the CD20$^+$ B cells for good outcome are more spread out as indicated by the higher value of the FD difference (Fig. 2C) and by the higher value of the area under the curve of the fraction of density hotspots vs. σ (Fig. 3C). In addition, good outcome patients have a higher density of B cells that infiltrate cancer cell islands (Fig. 4). This implies that when cells are more spatially dispersed, they are more likely to infiltrate cancer cell islands. We find support for this in Fig. 5A which shows a scatter plot for the correlation between the area under the hotspot curve vs. σ for CD20$^+$ B cells in the entire tumor tissue (including stroma and cancer cell islands) and the density of CD20$^+$ B cells in cancer cell islands. Each point represents a single patient. The correlation is significant for good outcome ($r = 0.54$, $p=0.007$) and for all patients ($r = 0.54$, $p=5 \times 10^{-4}$), but not for poor outcome ($r = 0.13$, $p=0.7$). Further support comes from Fig. 5B which shows the correlation between the FD difference for CD20$^+$ B cells in the entire tumor tissue and the density of CD20$^+$ B cells in cancer cell islands. Again, the correlation is significant for good outcome ($r = 0.51$, $p=0.01$) and for all patients ($r = 0.56$, $p=3 \times 10^{-4}$). (Analogous plots for CD8$^+$, CD3$^+$, CD4$^+$Foxp3$^-$, and Foxp3$^+$ T cells are shown in Fig. S10 where we see significant correlation between the hotspot AUC and the density of cells in cancer islands for CD8$^+$, CD3$^+$, and CD4$^+$Foxp3$^-$ T cells.)

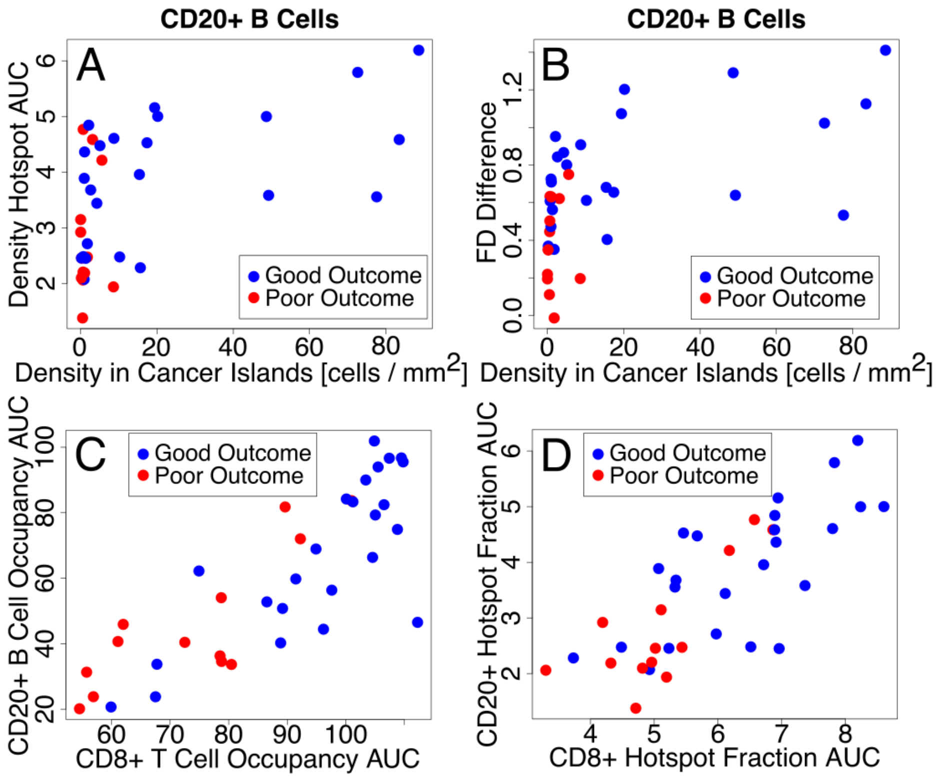

**Fig. 5: (A-B) Scatter plots showing correlations indicating that the more spread out cells are in the entire tumor tissue, the more likely they are to infiltrate cancer cell islands. (A) B cell hotspot AUC for entire tissue vs. B cell density in cancer cell islands. Pearson r = 0.54, p = 5 x $10^{-4}$. (B) B cell FD difference ([200-600 μm]- [10-40 μm]) for entire tissue vs. B cell density in cancer cell islands. Pearson r = 0.56, p = 3 x $10^{-4}$. (C-D) Scatter plots showing the correlation of spatial distributions of B and CD8 T cells in the entire tissue. (C) $CD20^+$ B cell occupancy AUC vs. $CD8^+$ T cell occupancy AUC in entire tissue. Pearson r = 0.83, p = 3 x $10^{-10}$. (D) $CD20^+$ B cell hotspot AUC vs. $CD8^+$ hotspot T cell AUC in entire tissue. Pearson r = 0.78, p = 2 x $10^{-8}$.**

10. Berthel A, *et al.* (2017) Detailed resolution analysis reveals spatial T cell heterogeneity in the invasive margin of colorectal cancer liver metastases associated with improved survival. *Oncoimmunology* 6(3):e1286436.
11. Saltz J, *et al.* (2018) Spatial Organization and Molecular Correlation of Tumor-Infiltrating Lymphocytes Using Deep Learning on Pathology Images. *Cell Rep* 23(1):181-193 e187.
12. Kather JN, *et al.* (2018) Topography of cancer-associated immune cells in human solid tumors. *Elife* 7.
13. Yuan Y (2016) Spatial Heterogeneity in the Tumor Microenvironment. *Cold Spring Harb Perspect Med* 6(8):1-18.
14. Maley CC, Koelble K, Natrajan R, Aktipis A, & Yuan Y (2015) An ecological measure of immune-cancer colocalization as a prognostic factor for breast cancer. *Breast Cancer Res* 17(1):131.
15. Getis A & Ord JK (1992) The Analysis of Spatial Association by Use of Distance Statistics. *Geographical Analysis* 24(3):189-206.
16. Nawaz S, Heindl A, Koelble K, & Yuan Y (2015) Beyond immune density: critical role of spatial heterogeneity in estrogen receptor-negative breast cancer. *Mod Pathol* 28(6):766-777.
17. Yuan Y (2015) Modelling the spatial heterogeneity and molecular correlates of lymphocytic infiltration in triple-negative breast cancer. *J R Soc Interface* 12(103).
18. Natrajan R, *et al.* (2016) Microenvironmental Heterogeneity Parallels Breast Cancer Progression: A Histology-Genomic Integration Analysis. *PLoS Med* 13(2):e1001961.
19. Mandelbrot BB (1983) *The Fractal Geometry of Nature* (W. H. Freeman and Co., New York) p 468.
20. Tambasco M, Eliasziw M, & Magliocco AM (2010) Morphologic complexity of epithelial architecture for predicting invasive breast cancer survival. *J Transl Med* 8:140.
21. Velanovich V (1998) Fractal analysis of mammographic lesions: a prospective, blinded trial. *Breast Cancer Res Treat* 49(3):245-249.
22. Chan A & Tuszynski JA (2016) Automatic prediction of tumour malignancy in breast cancer with fractal dimension. *R Soc Open Sci* 3(12):160558.
23. Baish JW & Jain RK (1998) Cancer, angiogenesis and fractals. *Nat Med* 4(9):984.
24. Baish JW & Jain RK (2000) Fractals and cancer. *Cancer Res* 60(14):3683-3688.
25. Lennon FE, *et al.* (2016) Unique fractal evaluation and therapeutic implications of mitochondrial morphology in malignant mesothelioma. *Sci Rep* 6:24578.
26. Bose P, *et al.* (2015) Fractal analysis of nuclear histology integrates tumor and stromal features into a single prognostic factor of the oral cancer microenvironment. *BMC Cancer* 15:409.
27. Bae MS, *et al.* (2016) Early Stage Triple-Negative Breast Cancer: Imaging and Clinical-Pathologic Factors Associated with Recurrence. *Radiology* 278(2):356-364.

28. Swets JA (1988) Measuring the accuracy of diagnostic systems. *Science* 240(4857):1285-1293.
29. Peitgen H-O, Jürgens H, & Saupe D (1992) *Chaos and fractals: new frontiers of science* (Springer-Verlag, New York).
30. Ali HR*, et al.* (2014) Association between CD8+T-cell infiltration and breast cancer survival in 12 439 patients. *Annals of Oncology* 25(8):1536-1543.
31. Liu S*, et al.* (2012) CD8+ lymphocyte infiltration is an independent favorable prognostic indicator in basal-like breast cancer. *Breast Cancer Res* 14(2):R48.
32. Li XF*, et al.* (2019) Infiltration of CD8(+) T cells into tumor cell clusters in triple-negative breast cancer. *Proceedings of the National Academy of Sciences of the United States of America* 116(9):3678-3687.
33. Binnewies M*, et al.* (2018) Understanding the tumor immune microenvironment (TIME) for effective therapy. *Nat Med* 24(5):541-550.
34. Boissonnas A, Fetler L, Zeelenberg IS, Hugues S, & Amigorena S (2007) In vivo imaging of cytotoxic T cell infiltration and elimination of a solid tumor. *J Exp Med* 204(2):345-356.
35. Kuczek DE*, et al.* (2019) Collagen density regulates the activity of tumor-infiltrating T cells. *J Immunother Cancer* 7(1):68.
36. Li X*, et al.* (2019) Infiltration of CD8(+) T cells into tumor cell clusters in triple-negative breast cancer. *Proc Natl Acad Sci U S A* 116(9):3678-3687.
37. Waliszewski P, Wagenlehner F, Gattenlohner S, & Weidner W (2015) On the Relationship Between Tumor Structure and Complexity of the Spatial Distribution of Cancer Cell Nuclei: A Fractal Geometrical Model of Prostate Carcinoma. *Prostate* 75(4):399-414.
38. Waliszewski P (2016) The Quantitative Criteria Based on the Fractal Dimensions, Entropy, and Lacunarity for the Spatial Distribution of Cancer Cell Nuclei Enable Identification of Low or High Aggressive Prostate Carcinomas. *Front Physiol* 7:34.
39. Baddeley A & Turner R (2005) spatstat: An R package for analyzing spatial point patterns. *Journal of Statistical Software* 12(6).
40. Pau G, Fuchs F, Sklyar O, Boutros M, & Huber W (2010) EBImage--an R package for image processing with applications to cellular phenotypes. *Bioinformatics* 26(7):979-981.

Supplementary Information for

### Fractal dimension, occupancy and hotspot analyses of B cell spatial distribution predict clinical outcome in breast cancer

Juliana C. Wortman[1#], Ting-Fang He[2#], Shawn Solomon[2], Robert Z. Zhang[2], Anthony Rosario[2], Roger Wang[2], Travis Y. Tu[2], Daniel Schmolze[3], Yuan Yuan[4], Susan E. Yost[4], Xuefei Li[5], Herbert Levine[5,6], Gurinder Atwal[7], Peter P. Lee[2+] and Clare C. Yu[1+]

[1]Department of Physics and Astronomy, University of California, Irvine, Irvine, CA 92697;
[2]Department of Immuno-Oncology, City of Hope Comprehensive Cancer Center and Beckman Research Institute, 1500 East Duarte Road, Duarte, CA 91010
[3]Department of Pathology, City of Hope Comprehensive Cancer Center, 1500 East Duarte Road, Duarte, CA 91010
[4]Department of Medical Oncology and Therapeutics Research, City of Hope Comprehensive Cancer Center, 1500 East Duarte Road, Duarte, CA 91010
[5]Department of Bioengineering and the Center for Theoretical Biological Physics, Rice University, Houston, TX 77030
[6]Department of Bioengineering and Department of Physics, Northeastern University, Boston, MA 02115
[7]Cold Spring Harbor Laboratory, Cold Spring Harbor, NY 11724
[#]Co-first authors

**This PDF file includes:**

Supplementary text
Table S1
Figs. S1 to S12

**Table S1: Clinicopathological characteristics of patients with TNBC. In the TNM classification, T denotes tumor size, N denotes lymph node invasion,**

**and M denotes the metastasis status. The patient samples were all used for antibody stains (N=37) except only some *() for CD4+Foxp3- (N=27).**

| | Good Outcome | Poor Outcome |
|---|---|---|
| Number of Patients | 24 *(19) | 13 *(8) |
| Mean Age | 55 *(56) | 58 *(57) |
| Age Range | 27 - 76 | 46 - 79 |
| Stage I | 5 *(5) | 7 *(3) |
| Stage II | 17 *(13) | 6 *(5) |
| Stage III | 2 *(1) | 0 *(0) |
| T1 | 7 *(7) | 7 *(3) |
| T2 | 16 *(11) | 6 *(5) |
| T3 | 1 *(1) | 0 *(0) |
| N0 | 19 *(15) | 11 *(6) |
| N1 | 3 *(1) | 2 *(0) |
| N2 | 2 *(2) | 0 *(0) |
| M0 | 3 *(0) | 0 *(0) |
| M1 | 0 *(0) | 0 *(0) |
| MX | 21 *(17) | 13 *(8) |
| Grade 1 | 0 *(0) | 0 *(0) |
| Grade 2 | 3 *(3) | 1 *(0) |
| Grade 3 | 21 *(16) | 12 *(8) |
| Mastectomy | 15 *(13) | 2 *(1) |
| Breast conserving surgery | 9 *(6) | 11 *(7) |

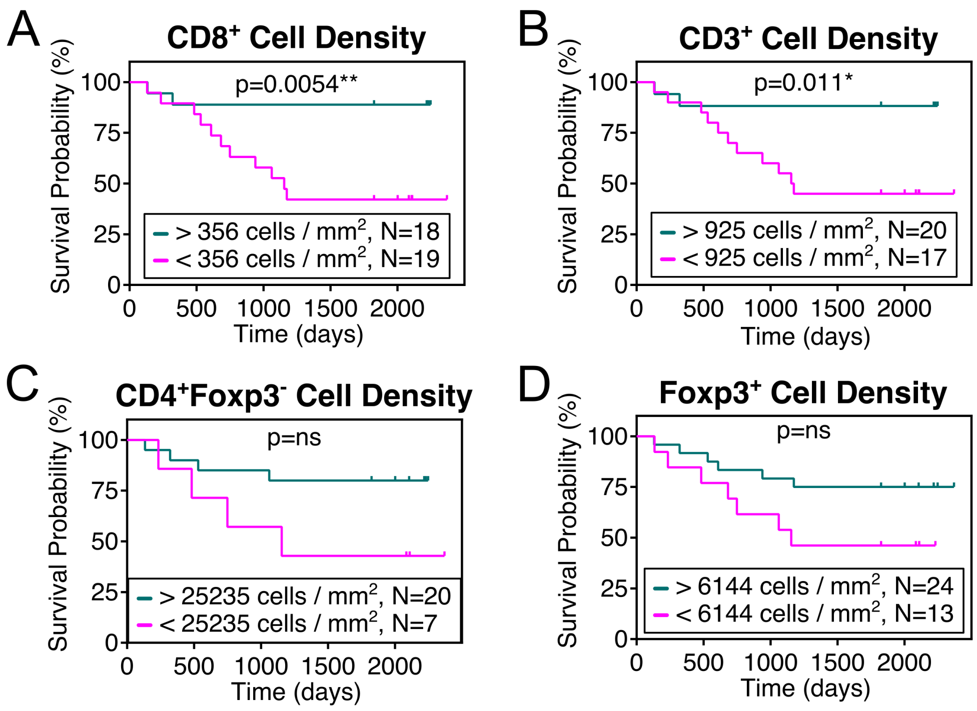

**Fig. S1: Relapse free survival (RFS) plots for (A) $CD8^+$ T cell density (B) $CD3^+$ T cell density (C) $CD4^+Foxp3^-$ Th cell density (D) $Foxp3^+$ Treg cell density. In the legends, N is the number of patients. Cell density is clinically significant for $CD8^+$ and $CD3^+$ T cells but not for Th and Tregs.**

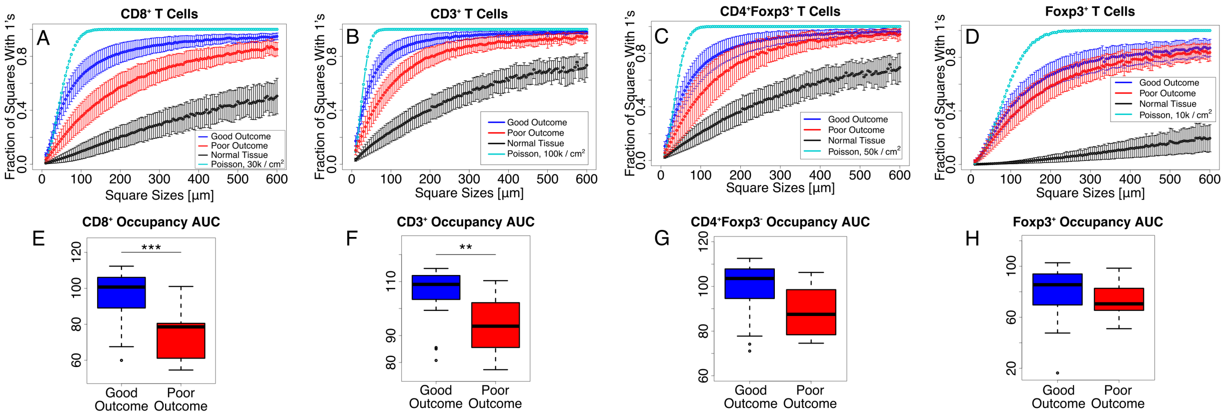

**Fig. S2: (A-D) Plots of occupancy vs. square size L for (A) CD8$^+$ T cells, (B) CD3$^+$ T cells, (C) CD4$^+$ Foxp3$^-$ T cells, and (D) Foxp3$^+$ T cells. (E-H) Box and whisker plots of occupancy AUC for (E) CD8$^+$ T cells, (F) CD3$^+$ T cells (G) CD4$^+$Foxp3$^-$ Th cells, and (H) Foxp3$^+$ Treg cells in the entire tissue. The occupancy AUC is clinically significant for CD8$^+$ and CD3$^+$ T cells but not for Th and Tregs.**

**Thinning:** In order to calculate the occupancy and fractal dimension, we overlay a grid of squares on the image and ask a binary question of each such as "Is there at least one CD20$^+$ B cell in the square?" (We will take B cells as a specific example, but the technique can be applied to any specific type of cell.) For questions such as this, the resulting occupancy or fractal dimension of B cells is somewhat dependent on the average density of B cells. To remove the dependence on B cell density, we "normalize" the cell density by a process we call "thinning". To do this, we look through the images from all the patients and find the image with the lowest average density of B cells. Then we randomly remove B cells in all the other images until all the images have the same B cell density. Here is a simple example of how to randomly remove B cells. To randomly remove half the B cells from an image, you would go to each B cell, flip a coin, and remove the cell if you get 'heads' and keep the cell if you get 'tails'. Fig. S3A illustrates thinning.

Figs. S3(B-K) show our results for thinned occupancy of CD20$^+$ B, CD8$^+$ T, and CD3$^+$ T cells. Note that the thinned CD20$^+$ B cell occupancy AUC is clinically very significant, much more so than the unthinned occupancy AUC for CD20$^+$ B cells (see Figs. 1C and 1E in main text). The thinned occupancy AUC for the T cells is either not clinically significant (CD8$^+$, Th, Tregs) or barely clinically significant (CD3$^+$ T cells).

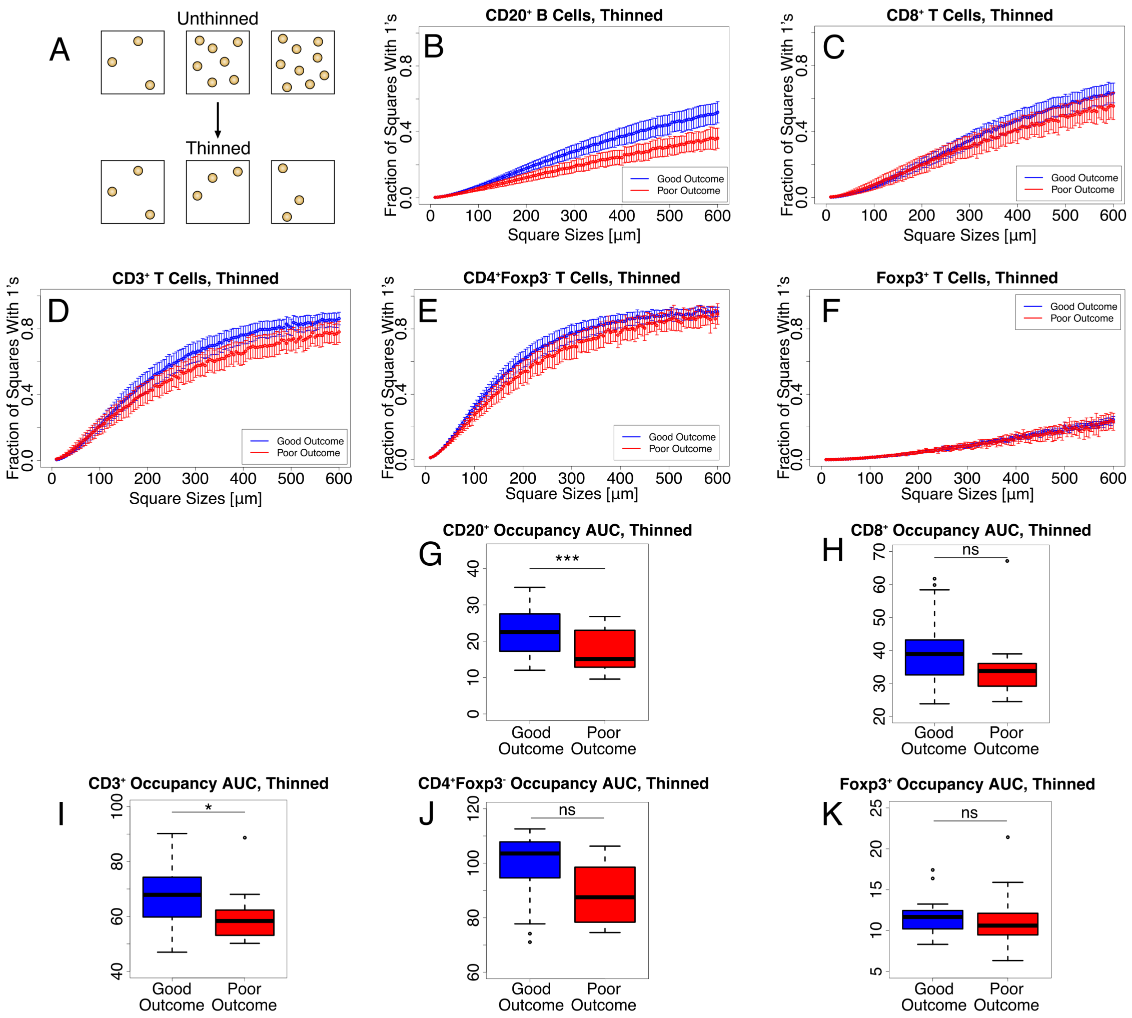

**Fig. S3: (A) Cartoon illustrating thinning of cell density. (B-F) Plots of occupancy vs. square size L for thinned cell densities. (B) CD20$^{+}$ B cells, thinned density = 12 cells / mm$^{2}$, (C) CD8$^{+}$ T cells, thinned density = 25 cells / mm$^{2}$, (D) CD3$^{+}$ T cells, thinned density = 120 cells / mm$^{2}$, (E) CD4$^{+}$Foxp3$^{-}$ Th cells, thinned density = 92 cells / mm$^{2}$, and (F) Foxp3$^{+}$ Treg cells, thinned density = 2.2 cells / mm$^{2}$. (G-K) Box and whisker plots of occupancy AUC for (G) thinned CD20$^{+}$ B cells, (H) thinned CD8$^{+}$ T cells, (I) thinned CD3$^{+}$ T cells, (J) thinned CD4$^{+}$Foxp3$^{-}$ Th cells, and (K) thinned Foxp3$^{+}$ Tregs in the entire tissue.**

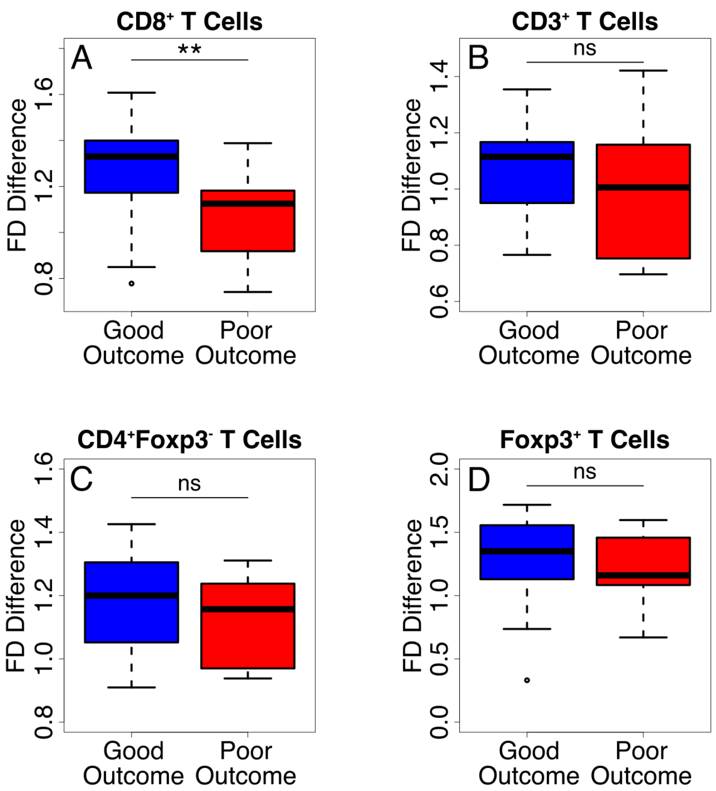

**Fig. S4: Fractal dimension (FD) difference: Box and whisker plots of FD difference (200 to 600 microns) – (10 to 40 microns) for unthinned (A) $CD8^+$ T cells (B) $CD3^+$ T cells (C) $CD4^+Foxp3^-$ Th cells (D) $Foxp3^+$ Treg cells. Note that the FD difference is correlated with clinical outcome only for $CD8^+$ T cells. For $CD8^+$ T cells, $\Delta s$ is larger for good outcome, indicating that the $CD8^+$ T cells are more spread out in this case.**

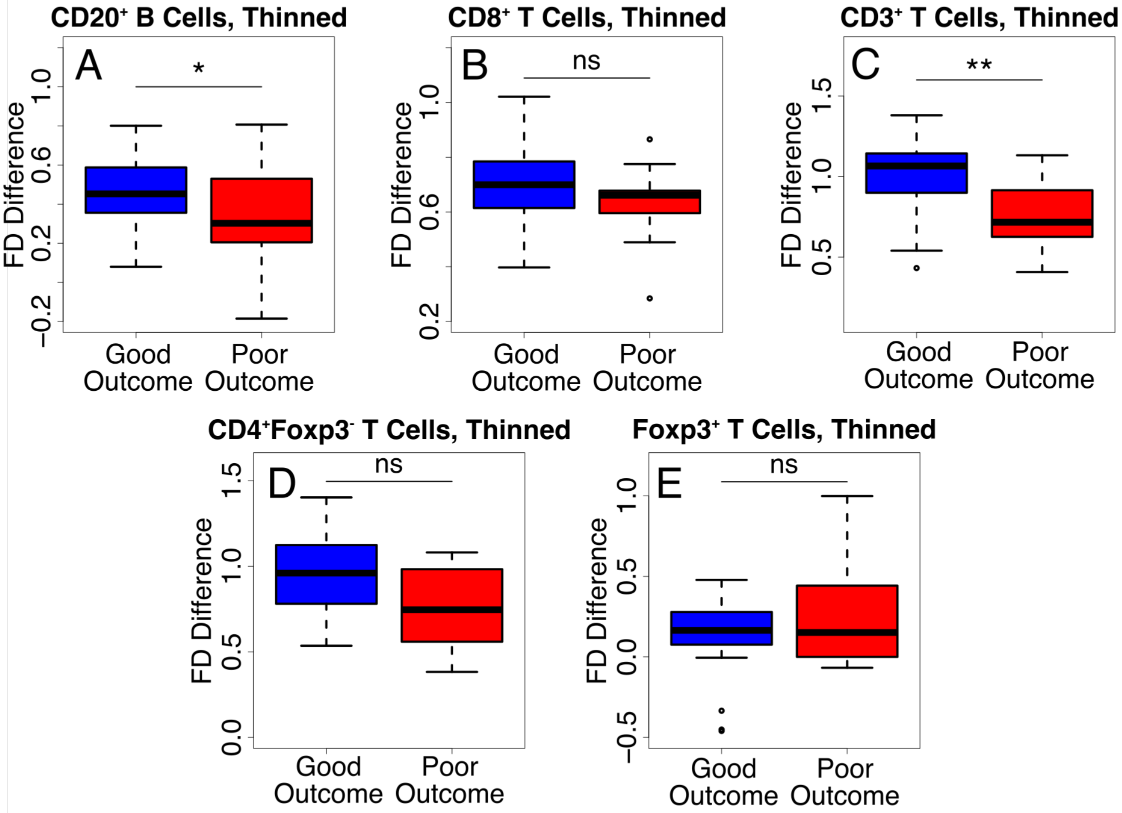

**Fig. S5: FD difference. Box and whisker plots of FD difference for thinned (A) $CD20^+$ B cells (12 cells/$mm^2$), (B) $CD8^+$ T cells (25 cells/$mm^2$), (C) $CD3^+$ T cells (120 cells/$mm^2$), (D) $CD4^+Foxp3^-$ T cells (92 cells/$mm^2$), and (E) $Foxp3^+$ T cells (2.2 cells/$mm^2$). Only thinned $CD20^+$ B cells and $CD3^+$ T cells have clinically significant FD differences. Note that for the thinned cells, we use a larger value for the short length scales because cells are farther apart compared to the unthinned ones. The FD difference is FD(200-600 microns) – FD(50-100 microns).**

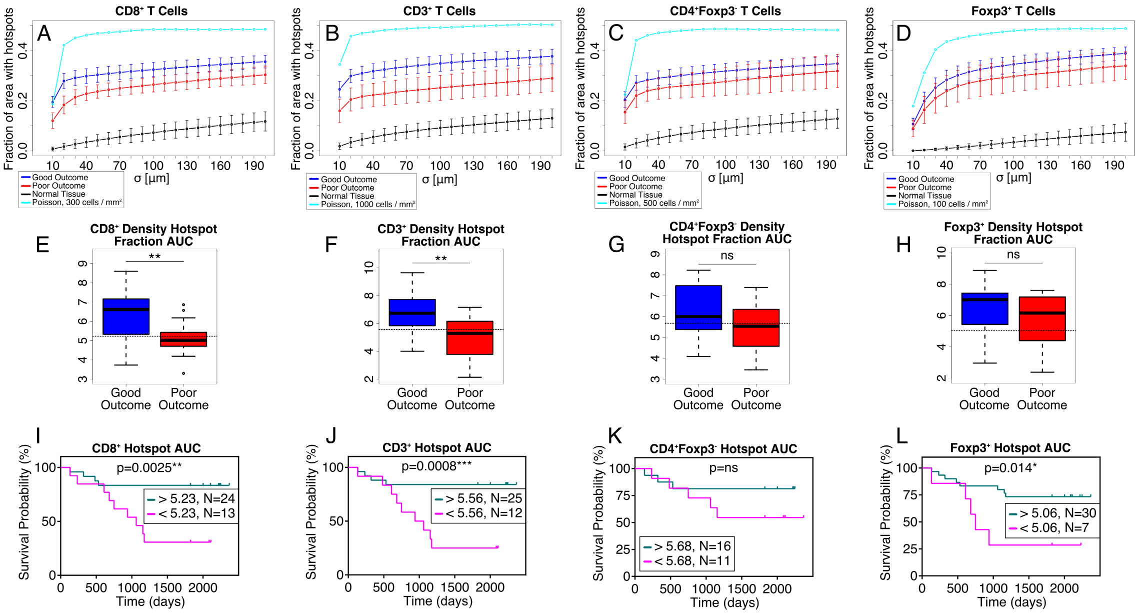

**Fig. S6: (A-D) Fraction of area with hotspots vs. σ for good clinical outcome (blue), poor clinical outcome (red), normal breast tissue (black) and randomly distributed points (cyan) for (A) $CD8^+$ T cells (B) $CD3^+$ T cells (C) $CD4^+Foxp3^-$ Th cells and (D) $Foxp3^+$ Treg cells. (E-H) Box and whisker plots of hotspot AUC for (E) CD8+ T cells (F) CD3+ T cells (G) $CD4^+Foxp3^-$ Th cells (H) $Foxp3^+$ Treg cells. (I-L) RFS plots for $CD8^+$ T cell hotspots (J) $CD3^+$ T cell hotspots (K) $CD4^+Foxp3^-$ Th cell hotspots and (L) $Foxp3^+$ Treg cell hotspots. Notice that the hotspot AUC is clinically significant for $CD8^+$ and $CD3^+$ T cells, but not for Th cells. The box and whisker plot for hotspot AUC for Tregs shows no significant difference for good and poor outcome (Fig. S6H), but the RFS plot shows a mildly significant difference. Note that in general, the patient cohorts of the high and low curves of the RFS plots are a mixture of what we have defined as good and poor clinical outcome.**

**Nearest neighbor (NN) distances between cells of a given type:** As we mention in the main text, one would think that a straightforward way to ascertain whether the B cells, say, are spread out spatially would be to determine the mean or median distance between nearest neighbor B cells (see Fig. S7A). However, most B cells are quite close (5-20 μm) to another B cell, so the NN distance just reflects the (inverse of the) local cell density rather than the spatial dispersion at long length scales. However, if the B cells (or cells of a given type) are thinned, then the mean or median nearest neighbor distances (post-thinning) can be a good measure of how spread out the cells are. The greater the post-thinning median or mean NN distance is, the more spatially dispersed the B cells are. Mean NN distances tend to be larger than median NN distances because a few isolated B cells that are far away from other B cells can skew the distribution and the average. We are just citing B cells as an example. The same considerations hold for T cells.

The data for NN mean and median distances are consistent with each other for a given cell type. To avoid the influence of isolated outlier cells, we will just present the median NN distances after thinning. Fig. S7B shows a box and whisker plot of the median distance between CD20$^+$ B cells. The results displayed in Fig. S7B are consistent with the findings in the main text that B cells are more spread out for good outcome. Similar results are shown for CD8$^+$ T cells (Fig. S7C) and CD3$^+$ T cells (Fig. S7D).

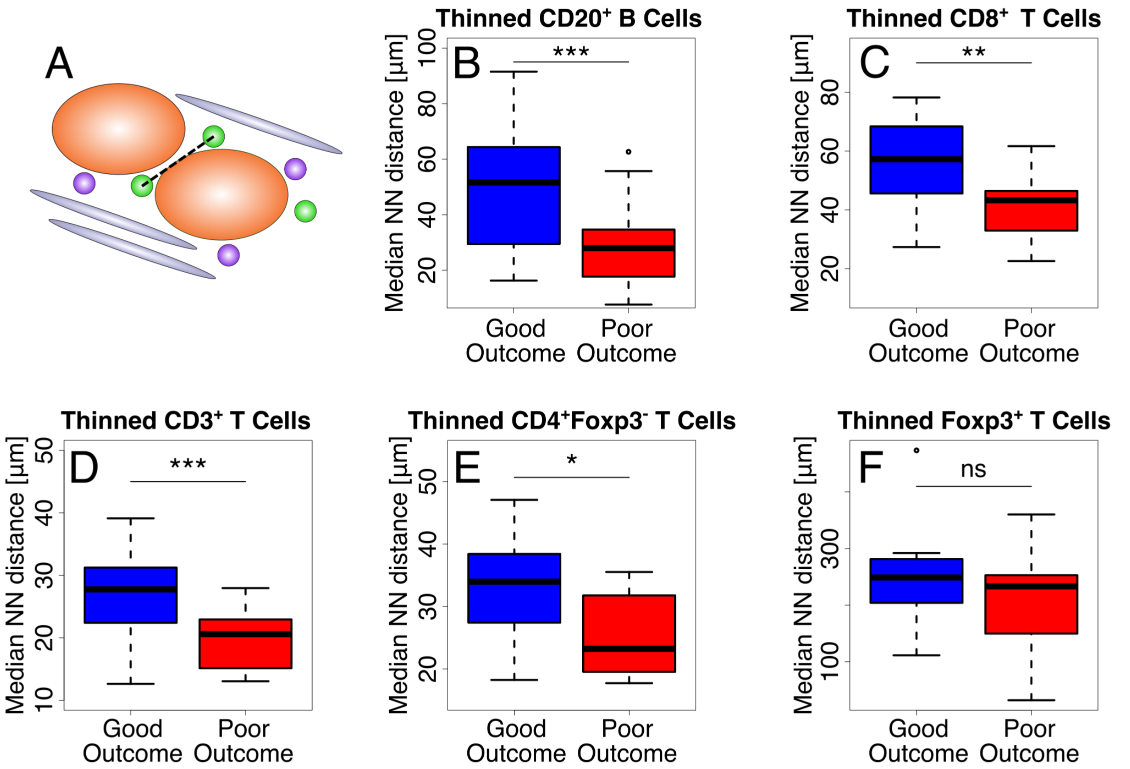

**Fig. S7: (A) Cartoon illustrating nearest neighbor (NN) distance between cells of the same type. (B-F) Box and whisker plots of the distribution of NN distances between thinned (B) CD20$^+$ B cells (C) CD8$^+$ T cells (D) CD3$^+$ T cells (E) CD4$^+$Foxp3$^-$ Th cells and (F) Foxp3$^+$ Treg cells. The difference in the thinned NN distributions between good and poor outcome shows a higher level of significance for CD8$^+$ T, CD3$^+$ T and CD20$^+$ B cells, and a lower level for CD4$^+$Foxp3$^-$ Th cells.**

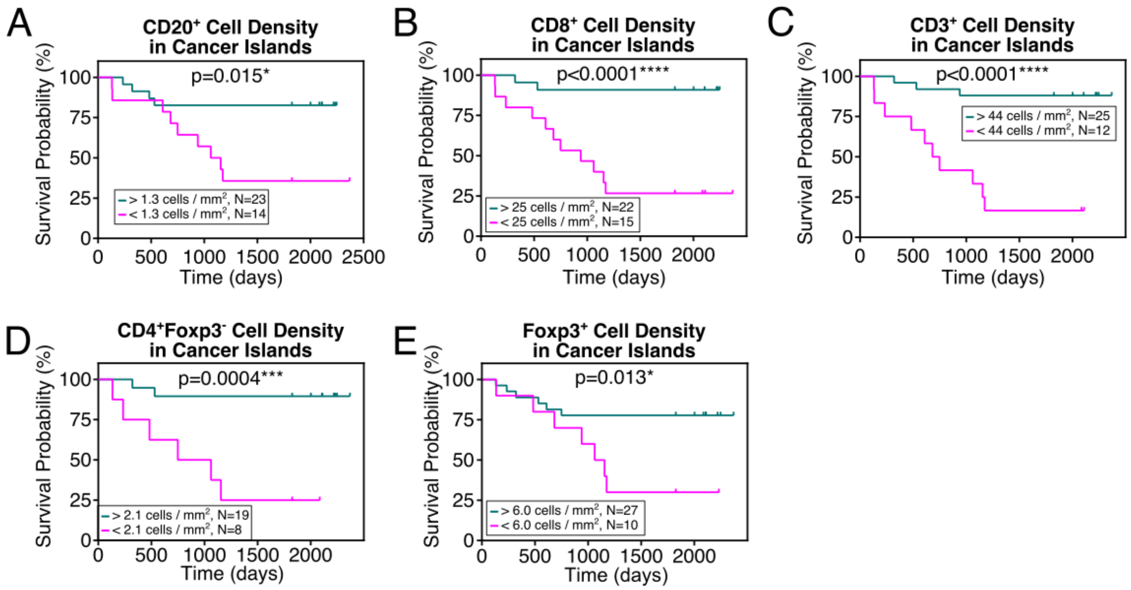

**Fig. S8: RFS plots of density of the following cells infiltrating into cancer cell islands: (A) $CD20^{+}$ B cells (B) $CD8^{+}$ T cells (C) $CD3^{+}$ T cells (D) $CD4^{+}Foxp3^{-}$ T cells and (E) $Foxp3^{+}$ T cells. Note that higher densities of B cells and all types of T cells in the cancer cell islands are correlated with improved prognosis.**

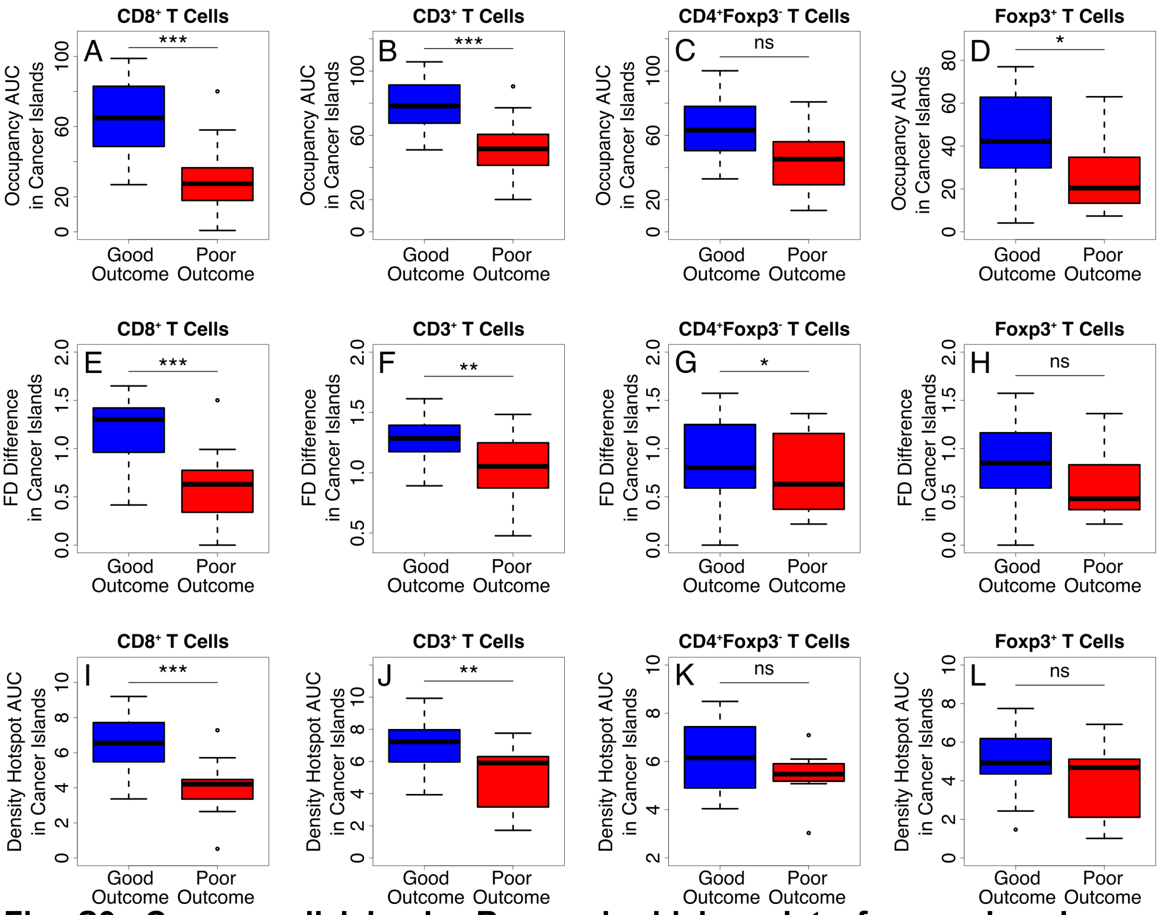

**Fig. S9: Cancer cell islands. Box and whisker plots for good and poor outcome for cells in cancer cell islands: (A) CD8$^+$ T cell occupancy AUC (B) CD3$^+$ T cell occupancy AUC (C) CD4$^+$Foxp3$^-$ T cell occupancy AUC (D) Foxp3$^+$T cell occupancy AUC (E) CD8$^+$ T cell FD difference [(200-600 microns) – (10-40 microns)] (F) CD3$^+$ T cell FD difference [(200-600 microns) – (10-40 microns)] (G) CD4$^+$Foxp3$^-$ T cell FD difference [(200-600 microns) – (10-40 microns)] (H) Foxp3$^+$ T cell FD difference [(200-600 microns) – (10-40 microns)] (I) CD8$^+$ T cell hotspot AUC (J) CD3$^+$ T cell hotspot AUC (K) CD4$^+$Foxp3$^-$ T cell hotspot AUC (L) Foxp3$^+$ T cell hotspot AUC. Note that the strong difference between the spatial distributions of CD8$^+$ and CD3$^+$ T cells for good and poor outcome is clinically significant.**

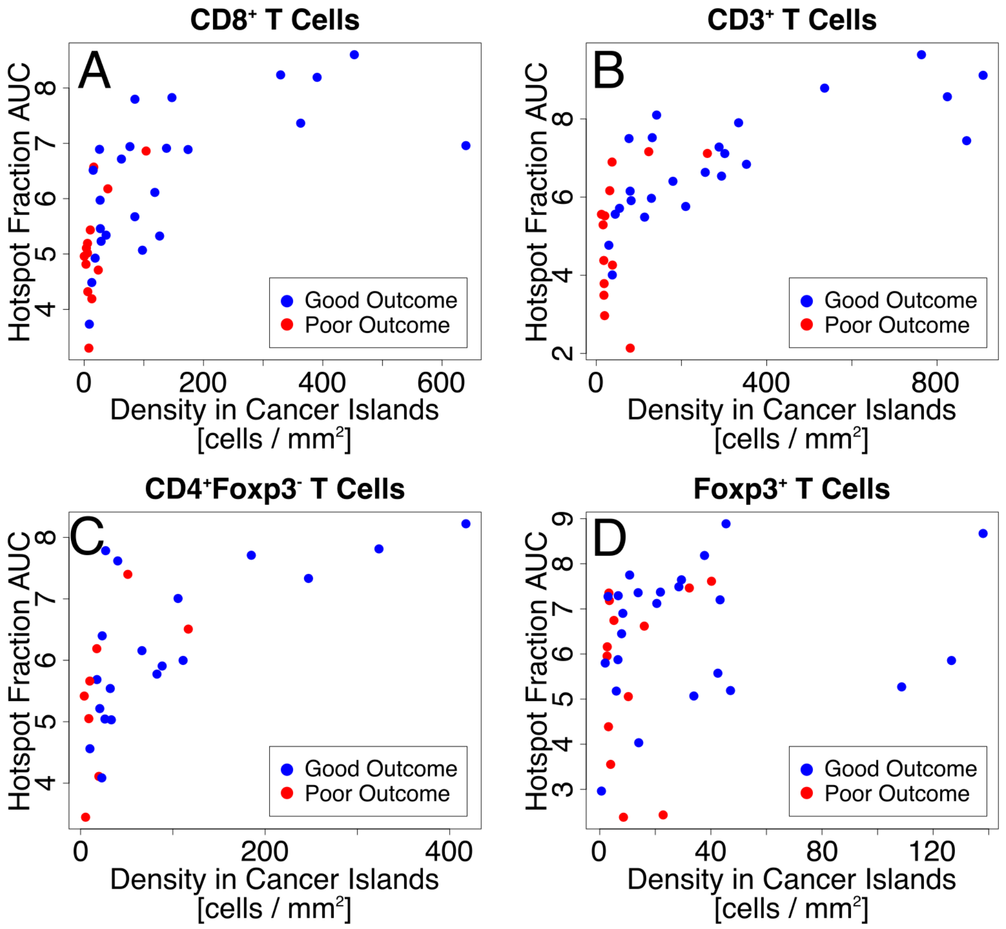

**Fig. S10: The more spread out cells are in the entire tissue, the more likely they are to infiltrate cancer cell islands. Scatter plots where each point represents a patient. (A) $CD8^+$ T cell hotspot AUC for entire tissue vs. $CD8^+$ T cell density in cancer cell islands. Pearson r = 0.66, p = 8.5 x $10^{-6}$. (B) $CD3^+$ T cell hotspot AUC for entire tissue vs. $CD3^+$ T cell density in cancer cell islands. Pearson r = 0.71, p = 8.63 x $10^{-7}$. (C) $CD4^+Foxp3^-$ T cell hotspot AUC for entire tissue vs. $CD4^+Foxp3^-$ T cell density in cancer cell islands. Pearson r = 0.66, p = 2.0 x $10^{-4}$. (D) $Foxp3^+$ T cell hotspot AUC for entire tissue vs. $Foxp3^+$ T cell density in cancer cell islands. Pearson r = 0.22, p = ns (0.20). Note the significant correlation for $CD8^+$, $CD3^+$, and $CD4^+Foxp3^-$ T cells.**

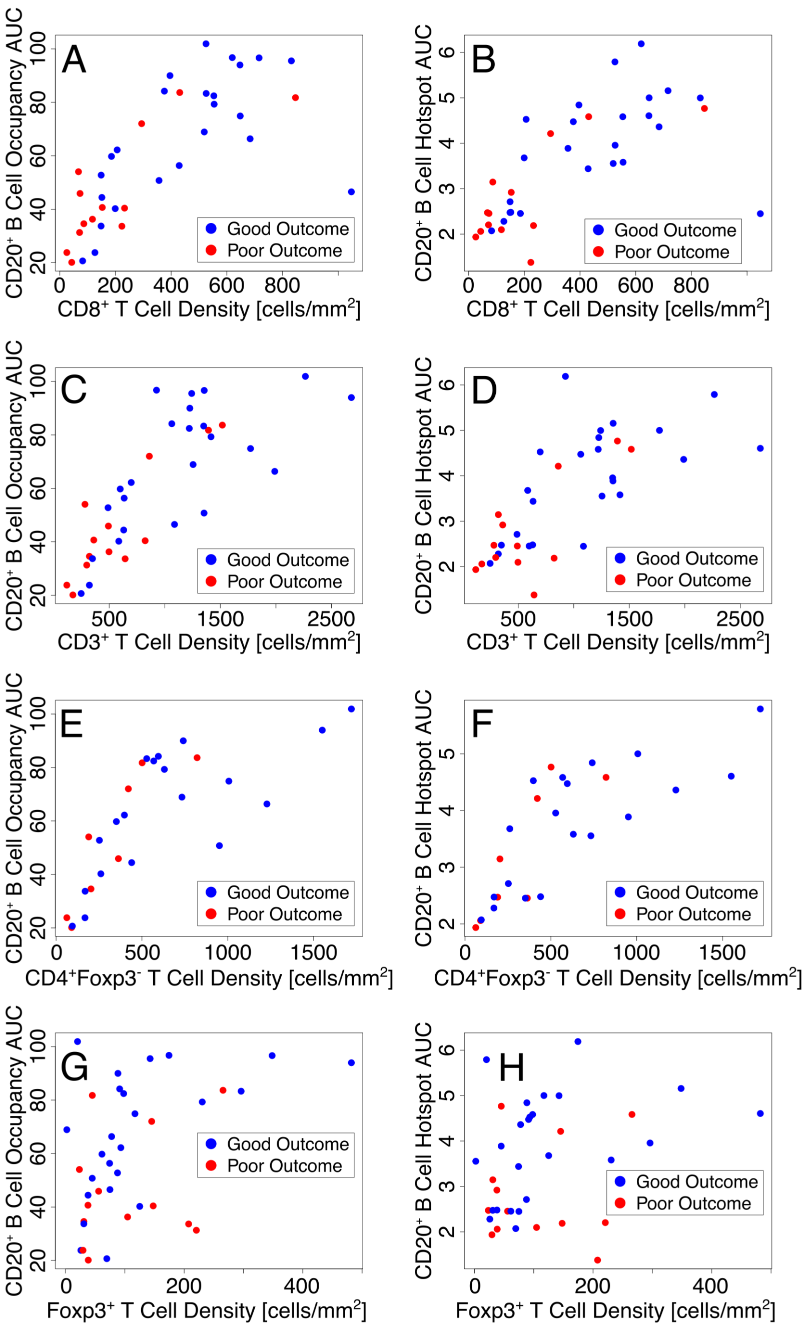

**Fig. S11: Scatter plots (entire tumor tissue) where each point represents a patient: (A) CD20$^+$ B cell occupancy AUC vs. the density of CD8$^+$ T cells. Pearson r = 0.72, p = 5.0 x 10$^{-7}$. (B) CD20$^+$ B cell hotspot AUC vs. the density of CD8$^+$ T cells. Pearson r = 0.66, p = 8.6 x 10$^{-6}$. (C) CD20$^+$ B cell occupancy AUC vs. the density of CD3$^+$ T cells. Pearson r = 0.79, p = 7.6 x 10$^{-9}$. (D) CD20$^+$ B cell hotspot AUC vs. the density of CD3$^+$ T cells. Pearson r = 0.72, p = 4.3 x 10$^{-7}$. (E) CD20$^+$ B cell occupancy AUC vs. the density of CD4$^+$Foxp3$^-$ T cells. Pearson r = 0.75, p = 7.2 x 10$^{-6}$. (F) CD20$^+$ B cell hotspot AUC vs. the density of CD4$^+$Foxp3$^-$ T cells. Pearson r = 0.77, p = 2.4 x 10$^{-6}$. (G) CD20$^+$ B cell occupancy AUC vs. the density of Foxp3$^+$ T cells. Pearson**

**$r = 0.44$, $p = 6.2 \times 10^{-3}$. (H) CD20$^{+}$ B cell hotspot AUC vs. the density of Foxp3$^{+}$ T cells. Pearson $r = 0.30$, $p$ = ns ($7.5 \times 10^{-1}$). This figure shows that the spatial distribution of CD20$^{+}$ B cells is strongly correlated with the density of CD3$^{+}$ T, CD8$^{+}$ T, and CD4+Foxp3- Th cells.**

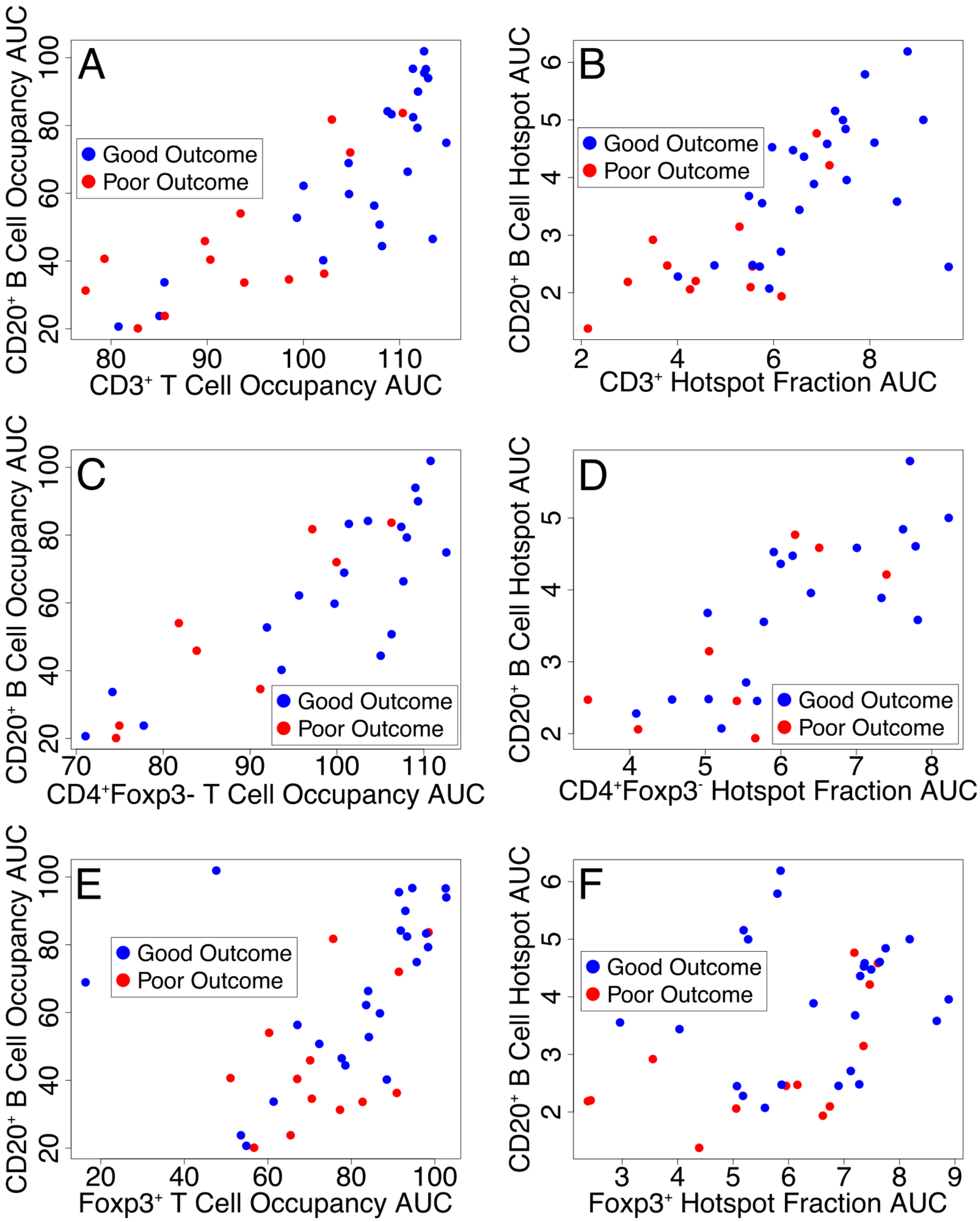

**Fig. S12: Scatter plot (entire tumor tissue): (A) CD20$^+$ B cell occupancy AUC vs. CD3$^+$ T cell occupancy AUC. Pearson r = 0.81, p = 1.4 x $10^{-9}$. (B) CD20$^+$ B cell hotspot AUC with CD3$^+$ T cell hotspot AUC. Pearson r = 0.70, p = 1.7 x $10^{-6}$. (C) CD20$^+$ B cell occupancy AUC with of CD4$^+$Foxp3$^-$ T cell occupancy AUC. Pearson r = 0.85, p = 2.6 x $10^{-8}$. (D) CD20$^+$ B cell hotspot AUC with of CD4$^+$Foxp3$^-$ T cell hotspot AUC. Pearson r = 0.77, p = 2.5 x $10^{-6}$. (E) CD20$^+$ B cell occupancy AUC with Foxp3$^+$ T cell occupancy AUC.**

**Pearson r = 0.47, p = 3.1 x $10^{-3}$. (F) $CD20^+$ B cell hotspot AUC with $Foxp3^+$ T cell hotspot AUC. Pearson r = 0.38, p = 2.0 x $10^{-2}$.**
